# Clinically-relevant treatment of PDX models reveals patterns of neuroblastoma chemoresistance

**DOI:** 10.1101/2022.04.01.486670

**Authors:** Adriana Mañas, Kristina Aaltonen, Natalie Andersson, Karin Hansson, Aleksandra Adamska, Alexandra Seger, Hiroaki Yasui, Hilda van den Bos, Katarzyna Radke, Javanshir Esfandyari, Madhura Satish Bhave, Jenny Karlsson, Diana Spierings, Floris Foijer, David Gisselsson, Daniel Bexell

## Abstract

Chemotherapy resistance and relapses are common in high-risk neuroblastoma (NB), an aggressive pediatric solid tumor of the sympathetic nervous system. Here, we developed a clinically-relevant in vivo treatment protocol mimicking the first line five-chemotherapy treatment regimen of high-risk NB and applied this protocol to mice with *MYCN*-amplified NB patient-derived xenografts (PDXs). Genomic and transcriptomic analyses were used to reveal the genetic and non-genetic mechanisms involved in NB chemoresistance. We observed convergent and parallel evolution of key NB genetic aberrations over time. Intrinsic resistance to chemotherapy was associated with high genetic diversity and an embryonic phenotype. Relapsed NB PDX tumors with acquired resistance showed an immature mesenchymal-like phenotype resembling multipotent Schwann cell precursors that are found in the adrenal gland. NBs with a successful treatment response presented a lineage-committed adrenergic phenotype similar to normal neuroblasts, reduced cell cycle gene expression, and negative regulation of the mitogen-activated protein kinase (MAPK)/extracellular signal-regulated kinase (ERK) cascade. NB organoids established from relapsed PDX tumors retained drug resistance, tumorigenicity, and transcriptional cell states ex vivo. This work sheds light on mechanisms involved in NB chemotherapy response in vivo and ex vivo using a clinically-relevant protocol, and emphasizes the importance of transcriptional cell states in treatment response. Detailed characterization of resistance mechanisms is essential for the development of novel treatment strategies in non-responsive or relapsed high-risk NB.

**One Sentence Summary:** COJEC chemotherapy treatment of neuroblastoma PDX models uncovers patterns of transcriptional plasticity and chemoresistance.

## INTRODUCTION

Neuroblastoma (NB) is a pediatric tumor that arises from the developing sympathetic nervous system and accounts for over 15% of childhood cancer deaths (*1*). Patients with high-risk tumors receive a very intense multimodal treatment regime that includes high doses of chemotherapy, surgery, radiation, stem cell transplants, and, in some cases, immunotherapy and targeted therapies. An established regimen to treat high-risk NB is the one set by the International Society of Pediatric Oncology – European Neuroblastoma (SIOPEN) (*2, 3*). Rapid COJEC is the induction chemotherapy step established by SIOPEN (*3, 4*). COJEC consists of high doses of five chemotherapeutic drugs (cisplatin, carboplatin, cyclophosphamide, etoposide, and vincristine) distributed in three courses that are alternatively administered in eight 10-day cycles. However, resistance is common. Some high-risk NB patients display upfront progression despite therapy; more commonly, patients initially respond but subsequently relapse, developing treatment-resistant disease. Thus, the overall survival for patients with high-risk disease remains lower than 50% (*5*).

Treatment resistance, which is a major clinical challenge and cause of death, is often linked to various aspects of intratumor heterogeneity. In principle, treatment resistance can arise from clonal evolution and selection of tumor cell subclones or by non-genetic mechanisms including transcriptional reprogramming (*6, 7*). NB is mainly driven by large chromosomal aberrations and is characterized by a low number of somatic driver mutations, even after relapse, and high inter- and intratumor heterogeneity (*8, 9*). Transcriptional and epigenetic analyses revealed that NB can exhibit at least two phenotypic cellular states, which are commonly referred to as ADR and MES (*10–19*). Cells in the ADR state are lineage-committed sympathetic nor-adrenergic cells. Cells in the MES state are immature neural crest cell-like or undifferentiated mesenchymal-like cells. Both states have been detected in established conventional cell lines (*10–14*) and in patient samples (*13–19*). In cell culture studies, MES cells are implicated in NB treatment resistance (*10,12, 20, 21*). However, this association has been difficult to demonstrate beyond cell culture. The effects of treatment pressure on NB cell states are unclear because patient studies use mainly untreated samples obtained during diagnosis, and cells with the MES phenotype have not been conclusively identified in xenograft studies in vivo (*12*).

The characteristically low number of somatic driver mutations, the small number of patients, and the heterogeneity within and across patients have made it especially challenging to identify the mechanisms of treatment resistance in NB (*8, 9*). A limitation of most preclinical studies of chemotherapy resistance is the use of single agents, rather than the combination of multiple drugs as is done for NB patients’ treatment. Here, we sought to investigate treatment resistance and relapse in NB in a clinically-relevant in vivo setting. We developed and used a COJEC-like treatment protocol that included all five COJEC drugs to treat multiple NB patient-derived xenograft (PDX) models that resemble the genotype and phenotype of NB patient tumors (*22, 23*). With this system, we detailed the transcriptomic and genomic changes occurring during NB treatment and at relapse.

## RESULTS

### Establishment of a COJEC-like treatment protocol using NB PDX tumors

Using three NB PDX models derived from high-risk NB tumors (*22, 23*), each with 1p deletion, *MYCN* amplification and 17q gain (Fig. S1A), we modeled NB treatment resistance using a treatment paradigm based on COJEC. PDX1 was derived from a COJEC-refractory tumor, whereas PDX2 and PDX3 were derived from COJEC-responsive tumors that subsequently relapsed (Fig. S1A). In our COJEC-like protocol, cisplatin, vincristine, etoposide, cyclophosphamide, and carboplatin were administered intraperitoneally in a cycled manner, for six 7-day cycles (Fig. S1B). Two tiers of dosing were designed and tested. The basic protocol (COJEC) was the lower dose and was well tolerated overall, whereas the high dose protocol (COJEC-HD) induced weight loss that required treatment interruptions to allow mice to recover (Fig. S1C). Both nude and NSG mice were evaluated and no difference in treatment response was observed between the mouse strains (Table S1). Mice were randomized into treatment groups when subcutaneous tumors reached ~500 mm^3^. Mice were treated intraperitoneally for six weeks with either saline (control), cisplatin (4 mg/kg, 3 times/week) or COJEC (Fig. 1A and S1B).

**Fig. 1.**
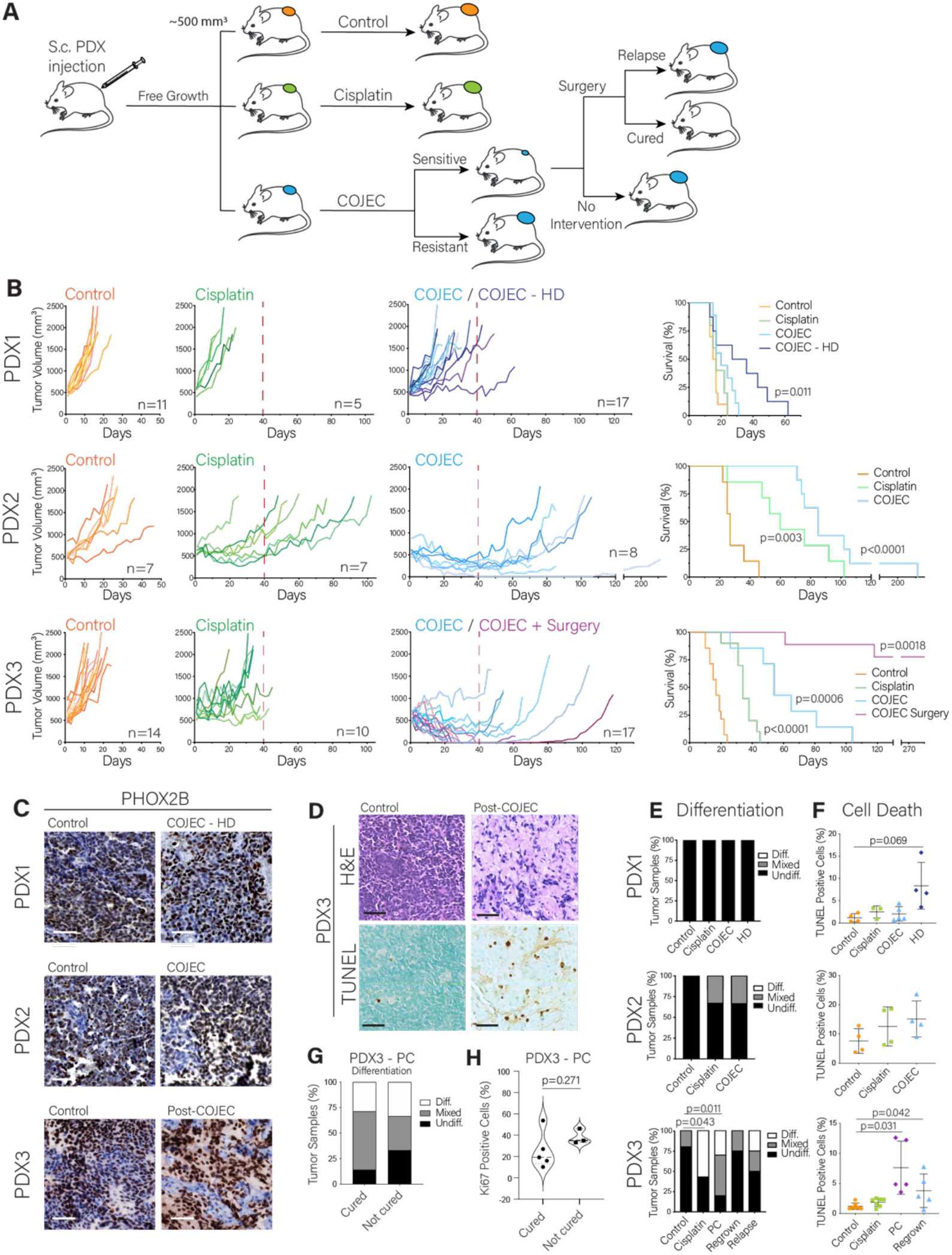
Establishment and application of a COJEC-like treatment protocol using NB PDX models. (**A**) Schematic representation of the experimental design. Mice were injected subcutaneously (S.c.) with dissociated PDX-derived organoids from PDX1, 2, or 3. When tumors reached ~500 mm^3^, mice were randomized into groups: Control, Cisplatin, and COJEC. A subgroup of PDX3 COJEC-responsive mice was further subjected to tumor resection surgery. (**B**) Tumor volume and mouse survival for each PDX model for the different treatment protocols. Red dashed line represents the end of COJEC. HD, high dose group for PDX1. Statistical analysis of survival was performed with log-rank test. (**C**) Immunohistochemical staining of PHOX2B (NB marker). (**D**) Representative Hoechst and eosin (H&E) and TUNEL staining for PDX3. (**E**) Assessment of differentiation status based on morphology from H&E staining. (**F**) Quantitation of cell death from TUNEL assay. (**G-H**) Quantitation of morphological differentiation and Ki67-positive cells for samples that were or were not cured in PDX3 in the PC group. Scale bars, 50 μm.

Consistent with the patient response, PDX1 tumors showed no or limited response to treatment with either cisplatin or the basic COJEC protocol (Fig. 1B). Therefore, we subjected this PDX model to the COJEC-HD protocol (Fig. S1B). Both COJEC doses led to smaller tumor volumes than controls on day 13 (median survival day for control group) (Fig. S1D), but only COJEC-HD treatment showed an increase in mouse survival (Fig. 1B). Overall, 19/20 PDX1 mice treated with cisplatin, COJEC, or COJEC-HD presented progressive disease (PD), and only one mouse showed stable disease (SD) during treatment (Table S1). These findings thus mirror the (lack of) therapy response seen in the corresponding patient.

Mice bearing PDX2 tumors responded well to treatment, presenting significant tumor size reduction (Fig. S1D; p < 0.05 for each therapy) and significantly increased survival when treated either with cisplatin or COJEC (Fig. 1B, p < 0.0001). Most COJEC-treated tumors (5/8) displayed a partial response (PR) and a significantly smaller tumor volume during treatment (Table S1 and Fig. S1D, p = 0.0008), whereas 4/7 cisplatin-treated tumors showed PD (Table S1). One COJEC-treated PDX2 mouse presented with complete response (CR) under treatment and relapsed locally after a period without treatment (Fig. 1B, Table S1).

COJEC-treated PDX3 tumors presented an overall PR, with significantly reduced tumor volume during COJEC treatment (Fig. S1D, p < 0.0001) and significantly increased survival (Fig. 1B, p = 0.0006), consistent with the corresponding patient’s clinical response (Fig. S1A). Two mice presented a CR and relapsed after treatment was removed (Table S1). To resemble the clinical regimen in patients, we performed surgical resection on a subgroup of COJEC-treated mice (n=9) when tumors had reduced below 200 mm^3^ (aimed volume reduction >60%). Following surgery, the mice did not receive any further COJEC treatment. At the end of the study, 2/9 mice in the COJEC + surgery group had relapsed locally (22% relapse rate) and 7/9 were “cured” (tumors did not relapse). Among the COJEC-treated PDX3 mice that did not have surgery, 2/7 mice relapsed after presenting a CR and 5/7 had tumors that exhibited a PR or SD but regrew rapidly after treatment cessation (Fig. 1B and Table S1). Among the 17 mice in the COJEC-treated group, one had to be euthanized early because of severe weight loss despite exhibiting SD with regard to the tumor. Because this mouse did not complete the 6-week COJEC protocol, for downstream analyses this tumor was grouped with the surgically collected samples in a group hereon referred as post-COJEC (PC), comprised of cured (n=7) and not-cured (n=3) PDX3 mice.

Thus, we developed and evaluated a COJEC-like protocol and showed that high-risk NB PDX models display clinically relevant and unique responses to this treatment, ranging from upfront PD to maintained CR. The results mirror those seen in the corresponding patients.

### Histological characterization of COJEC-treated NB PDX tumors

When subjected to immunohistochemical analysis, all tumors were positive for the NB marker PHOX2B (Fig. 1C) Hoechst and eosin (H&E) staining and blinded morphological analysis of tumor sections revealed that COJEC-treated PDX3 tumors (PC group) displayed clear signs of morphological differentiation (enrichment of neurofibrillary matrix), whereas this was less obvious in COJEC-treated PDX2 and absent in PDX1 tumors from mice receiving COJEC or COJEC-HD (Fig. 1D and E, Fig. S1E). TUNEL staining revealed increased cell death in PDX1 only for COJEC-HD treatment and in the PDX3 PC group (Fig. 1D and F, Fig. S1F). All tumors displayed intense staining for the proliferation marker Ki67 (Fig. S1G). PDX1 and PDX2 tumors exhibited large and collapsed blood vessels (Fig. S1G), consistent with a worse prognosis (*24*). PDX3 tumors had overall smaller and open blood vessels. Within the PDX3 PC group no statistically significant difference was seen in morphological differentiation or proliferation between the cured and non-cured samples (Fig. 1G and H).

### Clonal dynamics and signs of parallel and convergent evolution

To characterize the genetic changes occurring during and after COJEC treatment, we performed whole genome copy-number analysis of selected tumors within the control and treatment groups, as well as of parental tumor organoids before injection. A high number of subclones was detected in all three PDX models during tumor progression and treatment, and all models clearly showed a complex, branched evolutionary pattern (Fig. 2 and Fig. S2-S3). For each model, we defined a “stem” clone as having all copy number aberrations (CNAs) present in all organoids and tumors.

**Fig. 2.**
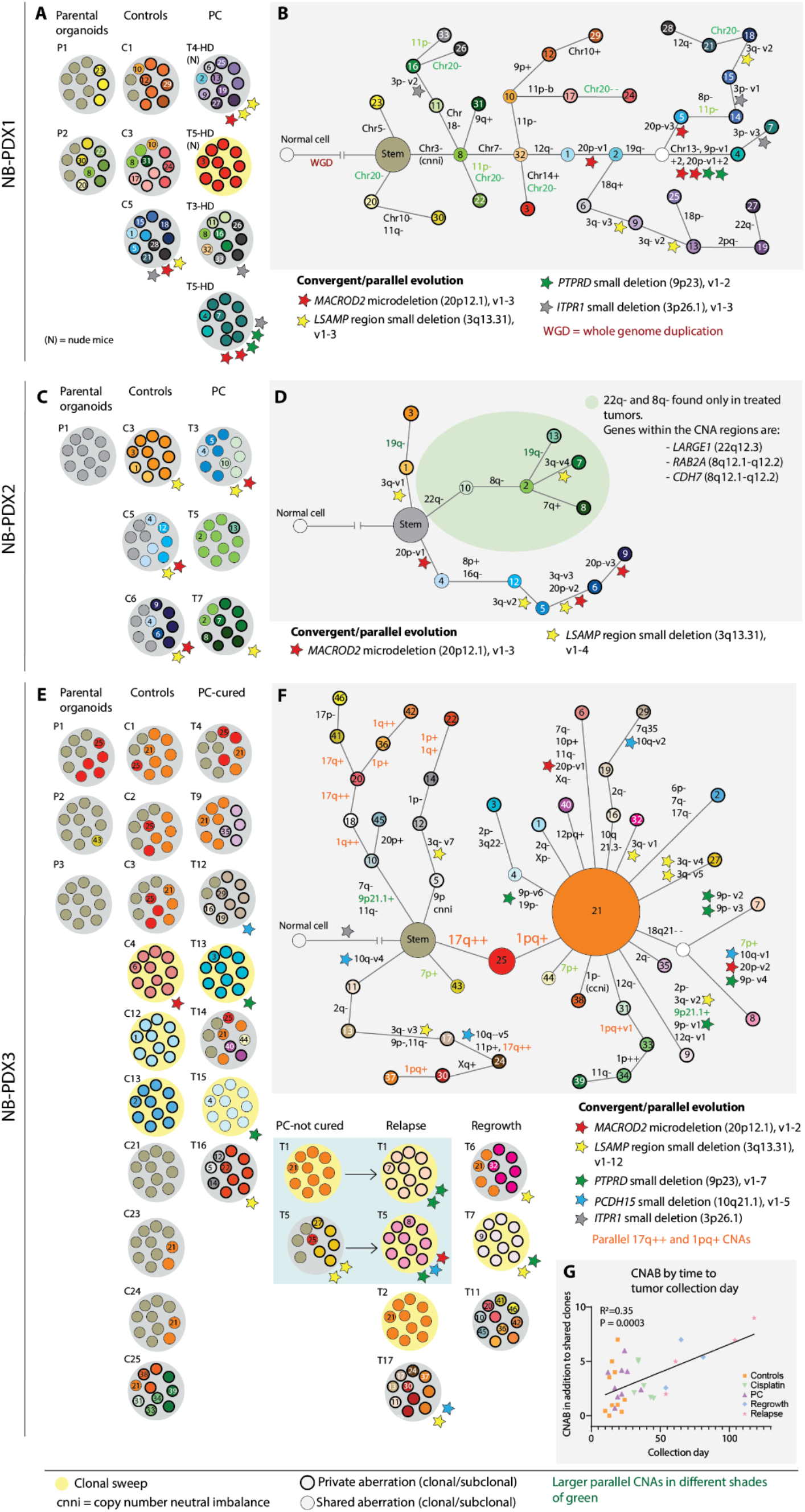
Genetic evolution of COJEC-treated NB PDXs based on copy number aberrations (CNAs). (**A-B**) PDX1, (**C-D**) PDX2, (**E-G**) PDX3. (**A, C, E**) Subclonal composition per sample where each small circle represents 10% of the tumor. Yellow background in the tumors illustrates a clonal sweep where one clone has taken over the tumor. Stars of different colors are signs of convergent and/or parallel evolution of specific genes (denoted for the respective tumor). Small circles with bold black outlines indicate private aberrations found only within that specific tumor. (**B, D, F**) Clone trees illustrating the phylogenetic relationship among the subclones, based on maximum likelihood phylogenetic analysis (see Fig. S2 and S3). (**G**) Correlation between CNAB and tumor collection day in PDX3. Statistical analysis by simple linear regression.

The non-responsive NB-PDX1 had a high number of subclones in tumors from both the control and the treatment groups (Fig. 2A). One tumor in a mouse that received COJEC-HD treatment exhibited a clonal sweep (Fig. 2A). A clone tree (Fig. 2B) based on phylogenetic analysis (Fig. S2A) illustrated how whole genome duplication (WGD) in the parental organoids created an abundance of genomic material and many subsequent whole chromosome or whole arm CNAs in all tumors. Complex branching and parallel evolution were detected, but no evidence of enrichment for specific subclones as a response to COJEC treatment was found.

For each NB-PDX, we summed the number of CNAs in each tumor or parental organoid to calculate the “copy number aberration burden” (CNAB). First, this was done for all aberrations detected in the respective stem as a comparison between the PDX models (Fig. S2B). After that, we calculated CNAB per tumor in addition to the phylogenetically defined stem clone (Fig. S2C). For PDX1, CNAB was comparable between controls and treated tumors (Fig. S2C), suggesting that COJEC-treatment did not increase the number of CNAs. However, the fraction of private aberrations (that is clones unique to a single tumor and representing high inter-tumor heterogeneity) was higher in the treated samples (Fig. 2A, Fig. S2D), suggesting diverse evolutionary trajectories in response to COJEC treatment in the individual tumors, despite genetically shared starting material. We analyzed the genetic diversity to describe subclonal variation within each tumor (Fig. S2E). Simpson’s Index of Diversity (Ds; 0 to 1 where 0 = no clonal variation and 1 = all subclones are different) showed no statistical difference between the controls and treated samples in PDX1 (Fig. S2E).

Compared to NB-PDX1, NB-PDX2 (Fig. 2C-D) had lower CNAB (Fig. S2C) and fewer private aberrations (Fig. 2D and S2D). Treatment-specific copy number losses were detected on chromosomes 22q and 8q (Fig. 2D, green shading, and Fig. S2F). The 22q region included a specific loss of a small part of *LARGE1* (encoding a glycosyltransferase) that was present in all treated samples. The 8q region included *RAB2A* (encoding a RAS family member involved in vesicular fusion and trafficking from the endoplasmic reticulum to the Golgi) and *CHD7* (encoding a DNA-binding protein with a chromodomain and helicase activity). CNAB calculation indicated that the total number of chromosomal aberrations was similar between controls and treated samples (Fig. S2C), as was genetic diversity (Fig. S2E). No clonal sweeps were detected in this model.

Most NB-PDX3 tumors showed a marked response to COJEC treatment. A high number of subclones and several selective sweeps occurred in both control tumors and treated samples (Fig. 2E-F, Fig. S3A-B). No specific subclone was selected by COJEC treatment, but we observed continuous clonal dynamics during tumor growth both in the presence and absence of treatment. These dynamics were evident by the trend of increased CNAB in the regrown and relapsed tumors (Fig. S2C), as well as in the linear correlation between high CNAB and time of tumor growth (Fig. 2G). Tumors from the control, cisplatin, and PC groups collected around the same time showed comparable CNAB (Fig. 2G). These data suggested that CNA events accumulate over time but that the number of different clones within tumors (genetic diversity) was not increased (Fig. S2E). We found two major CNA events occurring in the majority of the samples: an extra copy of the already gained q-region in chromosome 17 (17q++), clone 25 (Fig. 2E-F, red), and a new gain spanning part of the p-arm and all of the q-arm in chromosome 1 (1pq+), clone 21 (Fig. 2E-F, orange). These genomic events occurred in both controls and treated samples, and also in subclones that were phylogenetically not derived from clones 25 and 21 (Fig. 2F, orange text). The accumulation of the same genetic aberrations in tumors that were not clonally related represents a typical example of parallel evolution. Since the aberrations occurred in both controls and treated tumors it suggested that they resulted from tumor-intrinsic evolutionary forces rather than chemotherapy-induced selection.

There was evidence of parallel evolution of small deletions of specific genetic regions within each of the PDXs (colored stars, Fig. 2). Many of these genes were found in two or all three PDX models, suggesting convergent evolution in aggressive *MYCN*-positive NB. For example, deletions in *MACROD2* (20p12.1) and in *LSAMP* (3q13.31) occurred in controls and treated samples across the three PDXs; thus, are unlikely related to treatment. *MACROD2* microdeletions may cause chromosomal instability (CIN), and the gene could function as a tumor suppressor in CNA-driven tumors (*25*), but this is debated (*26*). In all three models, the *MACROD2* microdeletion was associated with extensive branching and the accumulation of CNAs (Fig 2B, D, F and Fig. S2A, F). *LSAMP* encodes a protein that is a neuronal surface glycoprotein and the gene has also been suggested to be a tumor suppressor (*27*). *PTPRD* (9p23) deletions were detected in PDX1 and 3. This gene has been suggested as a tumor suppressor in high-risk NB through destabilization of AURKA and MYCN (*28, 29*). Partial deletion of the gene *ITPR1* (3p26.1) was found in PDX1 (parallel evolution) and in PDX3 (stem). *ITPR1* encodes an intracellular calcium channel important for apoptosis in response to endoplasmic reticulum stress (*30*).

### High genetic diversity and clonal evolution over time revealed by single-cell DNA analysis

We performed single-cell DNA sequencing (scDNA-seq) and low pass single-cell copy-number aberration (CNA) analysis of selected PDX tumors to further dissect the subclonal composition and evolutionary patterns. scDNA-seq analysis of treatment-resistant PDX1 revealed a 58-81% of unique single-cell clones in each tumor (Fig. 3A, gray shades) with no clones detected in more than one tumor (Fig. 3A and Fig. S4A). No enrichment of distinct subclones after treatment could be found, consistent with the results from bulk DNA analysis. High genetic diversity (Fig. S4B) reflected a high number of cell-unique clones in control tumors as well as in tumors sampled post-COJEC. The WGD event, which we detected in the tumor-initiating organoids (Fig. 2B and Fig. S2A), provided high genetic diversity and genomic instability, which is seen in some patients (*31, 32*). Even though we found no indications that a specific clone could explain treatment resistance in this model, it is possible that a high adaptability caused by genomic instability contributes to an evolutionary mechanism for treatment resistance in PDX1 (Fig 3B).

**Fig. 3.**
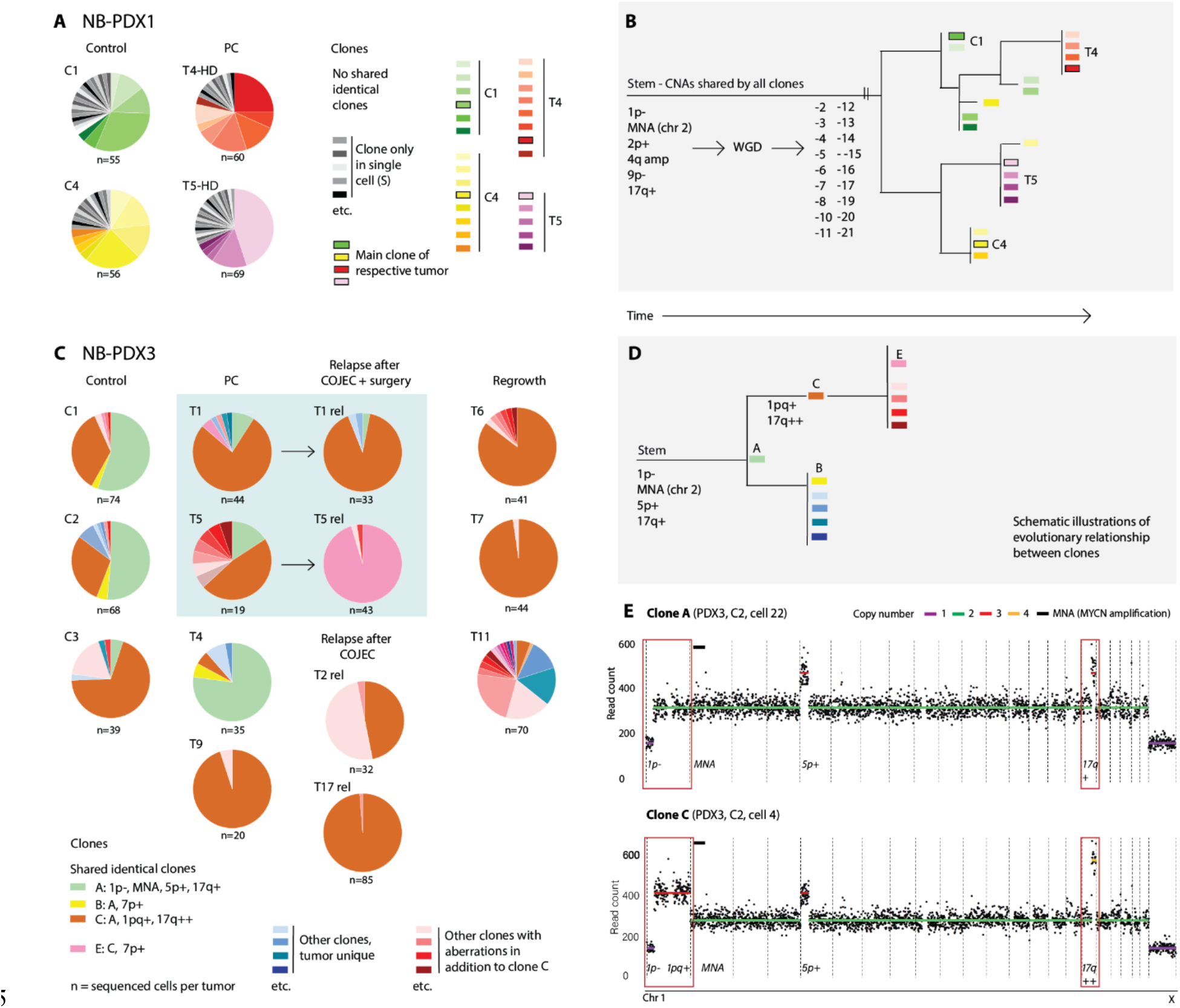
Clonal dynamics during and after COJEC treatment revealed by single-cell DNA sequencing. (**A**) Pie chart illustrating the fraction of different CNA-based clones within each of 2 control (C1 and C4) and 2 treated (T4-HD and T5-HD) NB-PDX1 tumors. Cells with a unique CNA from that in any other cell, that is a single-cell clone (S), are shown in shades of gray. No gray cell in any tumor is identical to a gray cell in another tumor from the same PDX. Clones represented by more than one cell are indicated in color. No shared identical clones means that, among these 4 NB-PDX1 tumors, each tumor had a unique set of clones. (**B**) Schematic view of the phylogenetic relationship among clones from the four NB-PDX1 tumors. WGD - whole genome duplication occurred in the stem and was subsequently followed by a high number of numeric losses shared by all clones. (**C**) Pie chart of clonal composition of NB-PDX3 tumors. Samples T1 and T5 were analyzed both at surgery (PC) and at a later relapse (rel). (**D**) Schematic view of the phylogenetic relationship between PDX3 clones A, B, C, and E. (**E**) Single-cell copy number profiles of clones A and C in PDX3. One profile per cell was received and processed in this analysis.

In the treatment-responsive PDX3, scDNA analysis revealed two major clones: clone A (1p-, MNA, 5p+, and 17q+) and clone C (1p-, 1pq+, MNA, 5p+, and 17q++) (Fig. 3C-D). Phylogenetic analysis showed that clone A (Fig. 3E) was more parental and that the aberrations associated with clone C (Fig. 3E, 1pq+ and 17q++) occurred after these changes (Fig. 3D and Fig. S4C). This result is consistent with data from bulk DNA analysis. The parental clone A was partly retained in control tumors and in tumors post-COJEC, but the occurrence of this clone was substantially decreased after regrowth and relapse, which represent tumors subjected to a treatment-induced genetic bottleneck (Fig. 3C). In contrast, clone C was enriched in tumors after treatment or at relapse (Fig. 3C, D) and showed aberrations commonly found in patient tumors (*33*). However, clone C was common in the controls and was not exclusive of treated tumors. Phylogenetic analysis indicated that, in addition to the dominating clones, unique NB cells can accumulate high numbers of CNAs (Fig. S4A and S4C). For example, two cells in PDX3 tumor T6 had gone through WGD, similar to PDX1. Enrichment of specific subclones did not lead to a significant decrease in genetic diversity in tumors after treatment and at relapse, but a trend was visible (Fig. S4B).

Taken together, scDNA-seq CNA analyses revealed contrasting patterns of clonal evolution between the treatment-responsive PDX3 and the refractory PDX1. PDX3 showed specific clonal enrichment of the clinically relevant 1pq and 17q regions. In contrast, PDX1 displayed an extremely high level of genetic diversity but no distinct clonal enrichment.

### Transcriptional signatures of COJEC treatment

We performed RNA-sequencing (RNA-seq) to explore the effects of COJEC at a transcriptional level. Unsupervised analysis of all the tumors (number of tumors: n=30 for PDX1, n=21 for PDX2, and n=41 for PDX3) showed that tumors strongly cluster within each PDX model (Fig. 4A, Fig. S5A and S5D). Analysis of each individual PDX revealed no distinct clusters for PDX1 (Fig. 4B) and PDX2 (Fig. 4C). However, for PDX3, a cluster representing PC-Cured tumors was observed (Fig. 4D). PDX1 tumors in nude mice or NSG mice had similar transcriptional profiles (Fig. S5B) and the treatment responses among PDX1 tumors from the two mouse strains was also similar (Fig S5C).

**Fig. 4.**
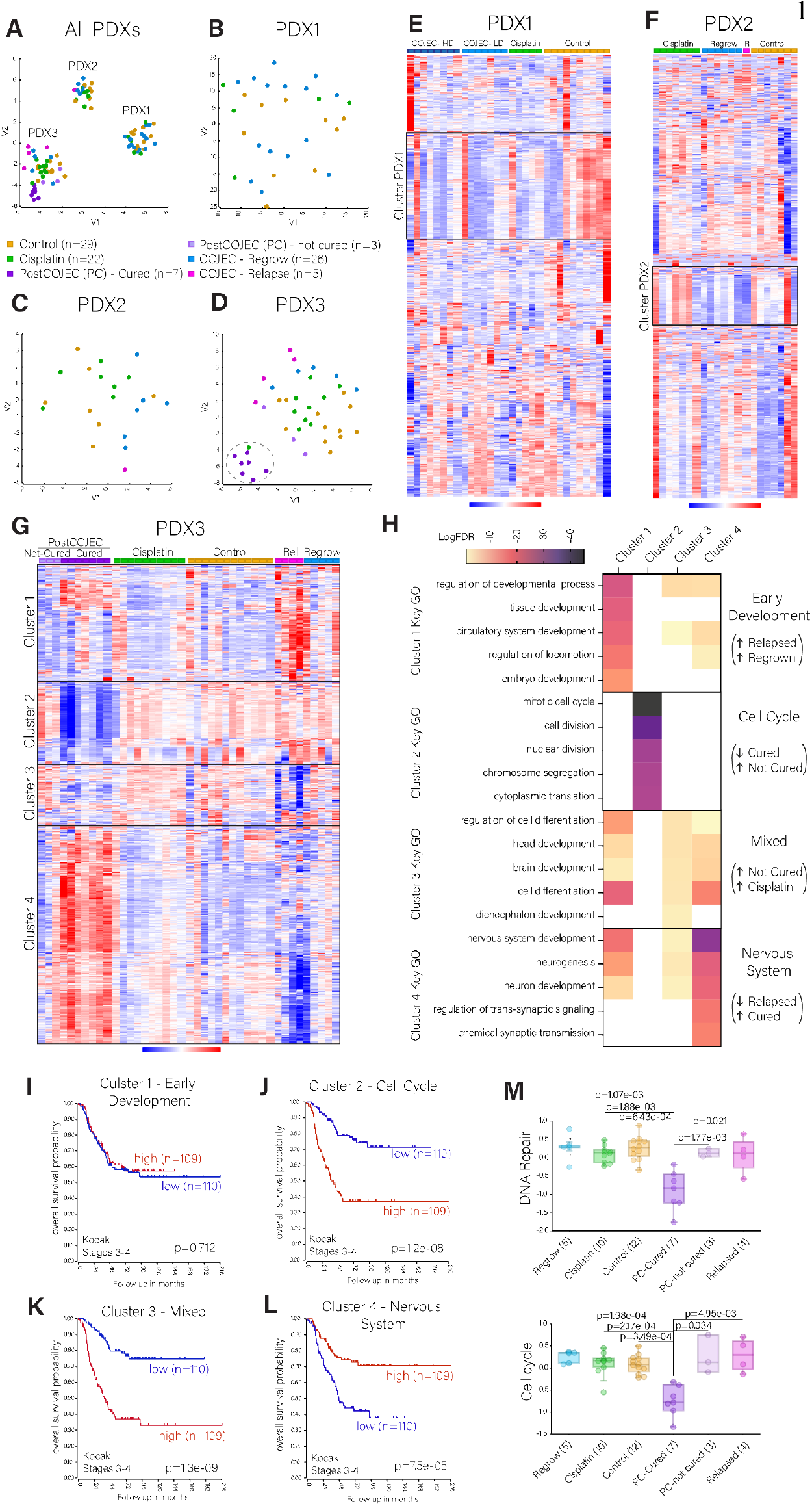
Transcriptomic analysis across NB PDXs and treatment groups. (**A-D**) t-SNE plots for all NB PDXs. Dotted line circle in (D) marks the cluster of PC-cured samples. (**E-G**) Unsupervised analysis of the top 1000 most variable genes across treatment groups for (E) PDX1, (F) PDX2, and (G) PDX3. Red is upregulated, blue is downregulated. Relapsed tumors (pink) are noted as R (for PDX2) and Rel (for PDX3). (**H**) Key gene ontologies (GO) defining each of the four gene clusters identified for PDX3 (G). (**I-L**) Overall survival Kaplan-Meier curves for each PDX3 gene cluster, identified from (G), in NB patients (stages 3 – 4, n=219; Kocak dataset, R2 Genomics Analysis Platform). (**M-N**) Expression (average z-score values over the gene set) of DNA repair and cell cycle gene signatures across PDX3 treatment groups. Statistical analysis performed with R2 genomics platform, ordinary one-way ANOVA with Welch’s t-test multiple comparison.

To establish a transcriptional baseline for each NB PDX, independent of treatment, we analyzed the top 1000 most variable genes expressed in the control tumors (Fig. S5E and Data file S1). Six well-defined clusters were identified. Gene ontology (GO) analysis revealed that PDX3 (Ctrl-Cluster 1) was mainly characterized by nervous system development genes and negative regulation of the MAPK/ERK cascade (Fig. S5F), whereas PDX1 (Ctrl-Cluster 6) contained mainly embryonic developmental signatures and a signature for regulation of Notch signaling (Fig. S5F). Few significant terms were identified for PDX2 with this analysis (Ctrl-Cluster 5). Analysis of differentially expressed genes (DEGs) with a false discovery rate (FDR) < 0.01 between control groups confirmed that PDX1 was characterized by upregulation of genes associated with embryonic development, the mitotic cell cycle, and metabolic processes (Fig. S5G-H). PDX2 had increased expression of genes associated with lipid metabolism along strong upregulation of the MAGE (melanoma associated gene) gene family (Fig. S5G, Fig. S5I). The MAGE gene family is linked to aggressive tumor progression and poor prognosis in several cancers (*34*). PDX3 was characterized by genes associated with nervous system development, differentiation, and regulation of ERK cascade (Fig. S5H, Fig. S5I). These results are consistent with our previous findings from the respective NB PDXs during *in vivo* passaging (*22*).

Next, we analyzed each PDX model to identify how COJEC treatment leads to transcriptional changes in NB. Unsupervised analysis revealed no clear transcriptional pattern in response to treatment for PDX1 and PDX2 (Fig. 4B and 4C, 4E and 4F). Analysis of the top 1000 most variable genes showed downregulation of one gene cluster in the COJEC groups for PDX1 and PDX2 (Fig. 4E and 4F, and Data file S1). In both cases, the downregulated gene clusters were mainly characterized by protein translation and protein localization ontologies (Fig. S6A and S6B). Only PDX3 samples showed a tendency to cluster by treatment, especially the cured samples and those that relapsed (Fig. 4D).

Unsupervised analysis of the top 1000 most variable genes in PDX3 revealed four gene clusters (Fig. 4G, and Data file S1). Using GO enrichment analysis, we characterized the clusters as “early development” (Cluster 1), “cell cycle” (Cluster 2), “mixed” (Cluster 3), and “nervous system” (Cluster 4) (Fig. 4H, Fig. S6C-S6F, and Data file S2). Relapsed and regrown tumors were characterized by increased expression of genes in early developmental pathways (Cluster 1) and downregulation of genes in nervous system development (Cluster 4) (Fig. 4G and 4H). A clear difference in expression was also observed between the surgery-collected samples that were cured (PC-cured) and those that later relapsed (PC-not cured), with the latter presenting an overall transcriptional signature similar to that of controls or cisplatin samples, with higher expression of clusters 2 and 3 (Fig. 4G). In contrast, the PC-cured tumors presented with strong downregulation of genes involved in the cell cycle (Cluster 2) and upregulation of nervous system development genes (Cluster 4) (Fig. 4G and 4H). This was further confirmed by DEG analysis of the PC-cured samples against the control, regrown, and relapse groups (Fig. S7A and Data file S1). The gene signatures defined by the PC-cured samples (low expression of Cluster 2 and high expression of Cluster 4) correlated with a significantly better prognosis in a cohort of 219 stages 3 – 4 NB patients (Fig. 4J and 4L, p < 0.0001). The mixed cluster 3, representative of the PC-not cured and cisplatin samples, correlated with poor prognosis in NB patients (Fig. 4K, p < 0.0001). PC-not cured tumors also showed significantly higher expression of DNA repair and cell cycle genes than the PC-cured samples (p=1.77e-03 and p=3.4e-02 respectively, Fig. 4M). Additionally, we explored the relevance of the MAPK/ERK and Notch signaling pathways, identified in the controls, and pertinent to many different cancers (*14, 35, 36*). The PC-cured samples presented significantly higher expression of genes involved in inhibition of the MAPK cascade (Fig. S7B, p < 0.01), and downregulation of Notch signaling (Fig. S7C, p < 0.05).

In summary, treatment-resistant and relapsed NB tumors exhibited features of early embryonic development. Tumors that eventually were cured showed features of nervous system development as well as reduced cell cycle and DNA repair gene expression following COJEC therapy, whereas tumors that eventually relapsed (represented by the PC-not cured samples) maintained a higher expression of cell cycle genes at the time of surgical removal.

### Identification of NB phenotypes based on transcriptional analysis

Our analysis revealed significantly enriched expression of nervous system development genes (Cluster 4) in PC-cured tumors (Fig. 5A, p < 0.01) and early development genes (Cluster 1) in relapsed tumors (Fig. 5B, p < 0.05). Previous studies identified ADR and immature MES-like tumor cell states in NB cell lines and patients tumors (*10–13, 15–19*), but their study using *in vivo* models has been challenging (*12, 20*) and the implications of chemotherapy are not clear. To explore these cell states, we analyzed the gene expression pattern of six pairs of publicly available ADR and MES-like signatures (*10, 12, 15–17, 19*) across the PDX3 treatment groups (Fig. S8A). Consistently, PC-cured tumors showed a higher ADR signature across all individual signatures. All of the immature MES-like signatures were higher in relapsed tumors, and some signatures were also high in PC-cured samples (Fig. S8A).

**Fig. 5.**
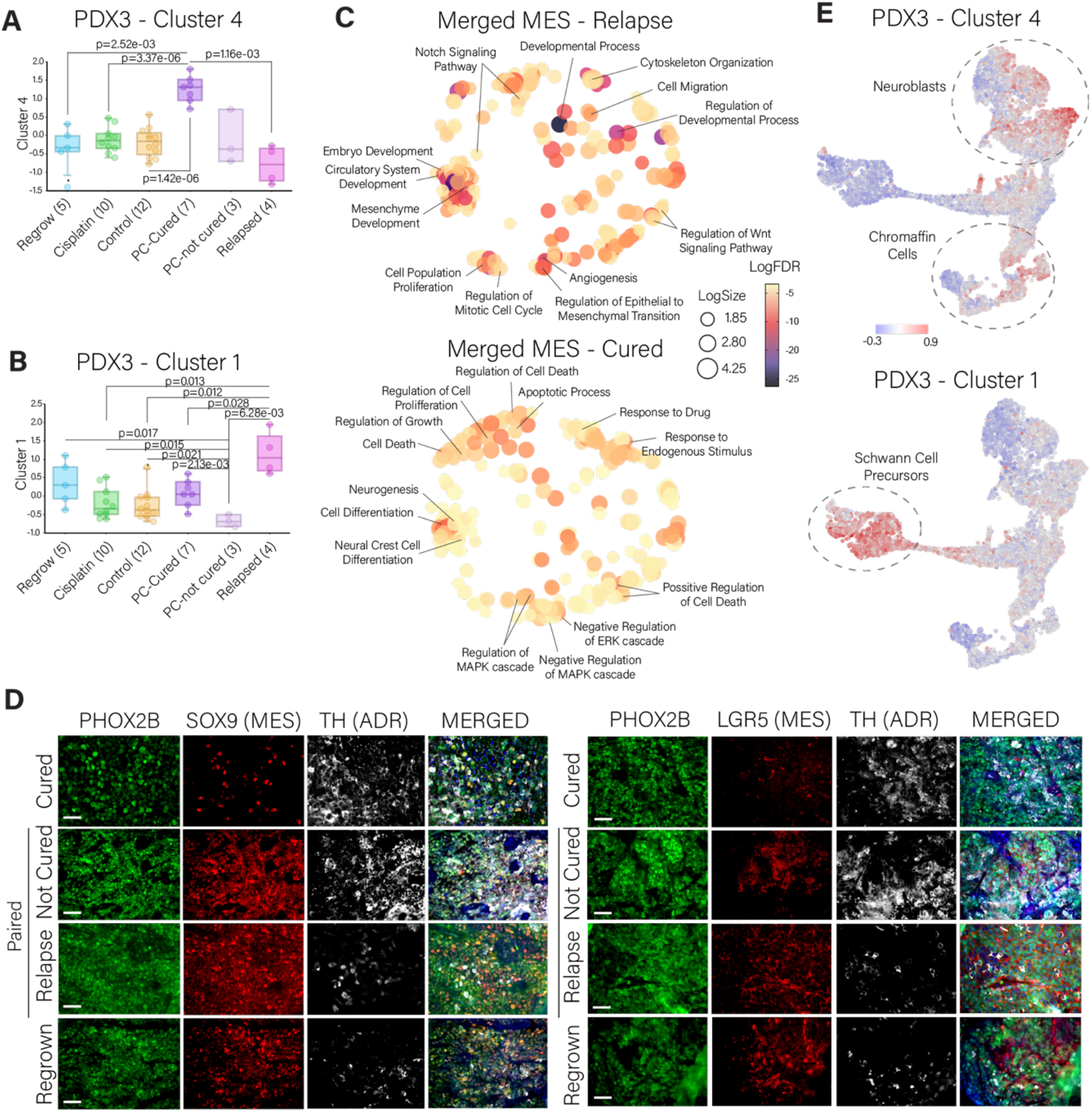
Adrenergic or mesenchymal-like gene signatures across NB PDX3 treatment groups. (**A, B**) Expression of PDX3-Cluster 4 (nervous system) and PDX3-Cluster 1 (early development) across PDX3 treatment groups. (**C**) Multidimensional scaling (MDS) analysis and visualization of the gene ontologies defined by the Merged-MES gene clusters corresponding to the relapsed and PC-cured tumors (Revigo, non-redundant scaling 0.7). (**D**) Immunofluorescence staining of PHOX2B (green), SOX9 (red, left panel), LGR5 (red, right panel), and TH (white) across PDX3 treatment groups. Blue nuclear staining with DAPI. Paired not cured and relapse samples correspond to different timepoints of the same mouse. Scale bars, 50 μm. (**E**) Mapping of PDX3-Cluster 4 and PDX3-Cluster 1 gene signatures over the Jansky et al. (*13*) single-cell RNA UMAP of normal adrenal medulla development.

Given the diverse definitions of “MES-like” or “immature” signatures in published work, we constructed two new gene signatures: “Merged ADR” is a signature comprised of our Cluster 4 genes and those ADR-associated genes in the 6 published signatures; “Merged MES” is a signature comprised of our Cluster 1 genes with all the MES or immature genes in the six pair published signatures. We analyzed the merged signatures across treatment groups for PDX3. Again, the ADR signature was enriched in the PC-cured tumors and the MES signature was enriched in the relapsed tumors (Fig. S8B). Similar to the association between low expression of cluster 4 with poor patient prognosis (Fig. 4L), low expression of Merged ADR correlated with poor prognosis in the same NB cohort (Fig. S8C). The Merged MES signature did not associate with NB patient prognosis (Fig. S8C). In summary, we observed that cured tumors display an enrichment of the ADR cell state, whereas the features of relapsed NB resembled the immature MES-like cell state.

### Transcriptional features of relapsed and cured NB

Based on the therapeutic responses, we explored in detail the ADR and MES gene expression signatures involved in relapsed, regrown, and cured samples by plotting the merged MES and ADR lists across these treatment groups (Fig. S8D and Data file S1). From the Merged MES list, we identified two sub-signatures: “MES-Relapse” (enriched in relapsed and regrown tumors) and “MES-Cured” (enriched in the PC-cured samples). GO analysis showed that the MES-relapse signature is enriched in early developmental pathways, cell proliferation, cell migration, and Notch and Wnt signaling (Fig. 5C). The MES-Cured signature is characterized by cell death, cell differentiation, and negative regulation of the MAPK/ERK cascade (Fig. 5C). From the Merged ADR list, we identified one main signature, Merged ADR-Cured (Fig. S8D).

MES-like signatures and markers obtained from NB patient tumors can be derived from NB cells or from stromal cells (*15, 16*). Here, we identified human-specific transcripts in the RNA-seq to distinguish human NB cells from mouse cells. Analysis of the fraction of RNA-seq “reads” specific to human and mouse showed that the small contribution of mouse reads was highly similar between PDX treatment groups (Fig. S9A-S9C). Only PC-cured samples in PDX3, which have a low MES signature, had a significantly higher presence of mouse stroma (Fig. S9C, p < 0.05). Relapsed samples had low percentages of mouse reads, indicating that the immature MES-like phenotype detected in relapsed tumors represents the NB cells rather than the stromal cells.

In the search for specific gene markers that correlate with prognosis both in the PDX model and in NB patients and have a minimal expression in stromal cells, we selected the common genes between the merged MES-Relapse and PDX3-Cluster 1 signatures, and the merged ADR-Cured and PDX3-Cluster 4 signatures, and filtered them through the cohort of 219 stage 3-4 patients (overall survival, median cut, FDR < 0.05). A curated list of 18 genes was identified from the MES lists and 146 genes from the ADR lists (Fig. S9D and S9E, and Data file S1), which correlated with poor and good prognosis in NB patients, respectively (Fig. S9F and S9G). Consistently, the 146-gene ADR signature was significantly higher in the PDX3 PC-cured samples, whereas the 18-gene MES signature was higher in relapse and regrown groups (Fig. S9H and S9I). Protein production from key genes selected from these lists was confirmed in PDX3 tumors (Fig. 5D). PHOX2B was used in our models as a pan-NB marker, similar to its clinical use. Consistent with the RNA results, cured tumors show an overall low protein expression of SOX9 and LGR5 (MES markers), and high abundance of TH (ADR marker), whereas relapse and regrown tumors showed the opposite pattern. Interestingly, PC-not cured samples present high amounts of both MES and ADR markers at the surgical resection time point (Fig. 5D), pointing to the importance of the immature MES-like cells in drug resistance and relapse *in vivo.*

### Relapsed NB resembles human Schwann cell precursors

NB is derived from progenitors of the sympathetic nervous system. We analyzed how the gene clusters 1 (relapsed tumors) and 4 (PC-cured) relate to the human adrenal medulla during normal development (*13*). Genes in the Cluster 1 signature were highly expressed in immature Schwann cell precursors (SCPs), whereas high expression of Cluster 4 genes mainly correlated with neuroblasts and chromaffin cells (Fig. 5E). Thus, COJEC-treated relapsed NB PDX tumors resemble human SCPs in the adrenal medulla, which is consistent with other reports (*16, 37, 38*).

### MYCN transcript and protein abundance in response to COJEC treatment

The *MYCN* gene is essential in NB pathogenesis (*39, 40*). Based on the scDNA data, we determined the number of *MYCN* copies for PDX1 and PDX3 (Fig. 6A and 6B). The non-responsive PDX1 had higher *MYCN* copy numbers compared to the responsive PDX3 (Fig. 6A). Within PDX3, all treatment groups had similar numbers of *MYCN* copies (Fig. 6B). We next analyzed *MYCN* RNA expression across PDX models and treatment groups (Fig. 6C). Consistent with CNAs, the amount of *MYCN* RNA was lower in PDX3 compared to PDX1 (Fig. 6D). PC-cured tumors presented strong *MYCN* RNA downregulation, whereas relapsed and PC-not cured tumors showed expression in the range of PDX1 and PDX2 (Fig. 6C and 6E). Consistent with the PDX1 and PDX2 tumors exhibiting higher *MYCN* transcripts, these tumors had high expression of a MYCN target gene signature (*41*) (Fig. 6F). PC-cured tumors also had lower expression of MYCN downstream target genes (Fig. 6G). At the protein level, we observed that relapse samples had a significantly higher number of MYCN-positive cells than PC-cured samples (Fig. 6H and 6I, p=3.8e-03). Thus, NB responding to COJEC showed downregulation of *MYCN* RNA and protein and its target genes, despite high *MYCN* copy numbers.

**Fig. 6.**
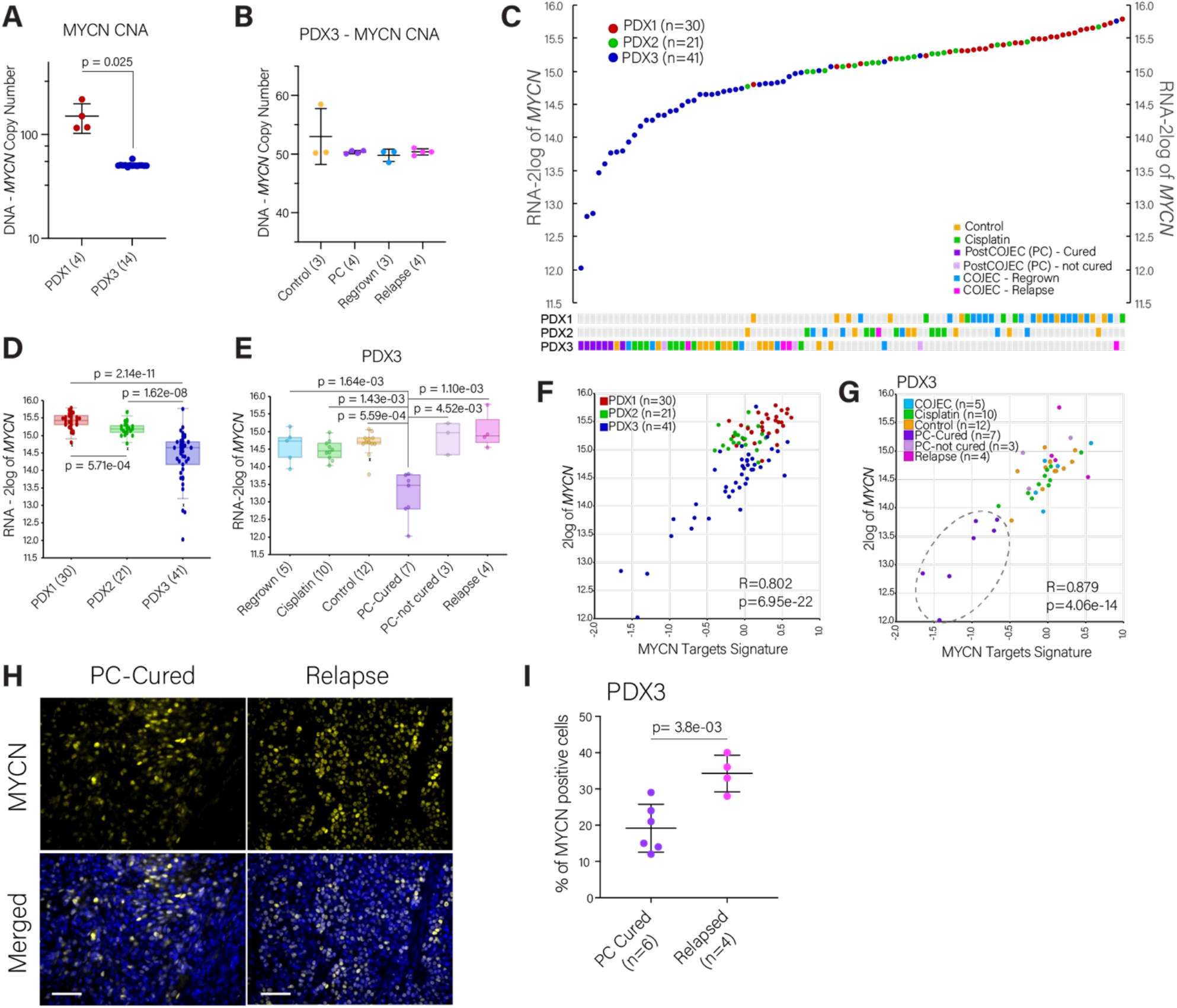
MYCN transcript and protein abundance during and after COJEC treatment. (**A**) Number of *MYCN* amplification copies in PDX1 and PDX3 tumors obtained from scDNA analysis. (**B**) Number of *MYCN* amplification copies across PDX3 treatment groups obtained from scDNA analysis. (**C**) *MYCN* gene expression for each individual tumor across all PDXs. (**D, E**) Statistical analysis of *MYCN* gene expression (transcripts) across NB PDXs (D) and across PDX3 treatment groups (E). (**F, G**) Correlation between *MYCN* expression and MYCN targets signature across PDXs (F) and PDX3 treatment groups (G). (**H**) Immunofluorescence staining of MYCN across PDX3 treatment groups. Blue nuclear staining with DAPI. Scale bars, 50 μm. (**I**) Statistical comparison by Welch’s t-test of MYCN staining quantitation from fluorescence staining. Symbols represent individual tumors.

### Influence of key genetic aberrations in transcriptional signatures and treatment response

Based on bulk DNA-seq data (Fig. 2), we observed parallel and convergent evolution of certain fragment losses that affected eight genes (Fig. S9J). These specific convergent losses seemed to accumulate across COJEC-treated and cisplatin-treated groups (Fig. S9J), but they did not correlate with treatment response. Instead, we found that the transcript abundance of these genes correlated with response: PC-cured samples had significantly higher expression of this 8-gene signature (Fig. S9J and S9K, p < 0.05). High expression of this signature also correlated with good prognosis in patients (Fig. S9L). Further work is needed to clarify the clinical relevance of these genes.

### 3D tumor organoids from relapsed NB maintain in vivo phenotype and tumorigenicity

We established free-floating 3D NB organoids, cultured under serum-free conditions with the addition of epidermal growth factor (EGF) and fibroblast growth factor 2 (FGF2), from control and COJEC-treated tumors for all three PDX models (Fig. 7A-7C). All NB organoids were tested with cisplatin and vincristine as single drugs to assess for acquired resistance to chemotherapy (Fig. S10A-S10C). NB organoids derived from PDX1 and PDX2 showed no difference in drug responses between control and COJEC-treated (Fig. S10A-S10B). However, NB organoids derived from a relapsed PDX3 tumor (tumor T5) showed decreased treatment response as compared to PDX3 control organoids (Fig. S10C).

**Fig. 7.**
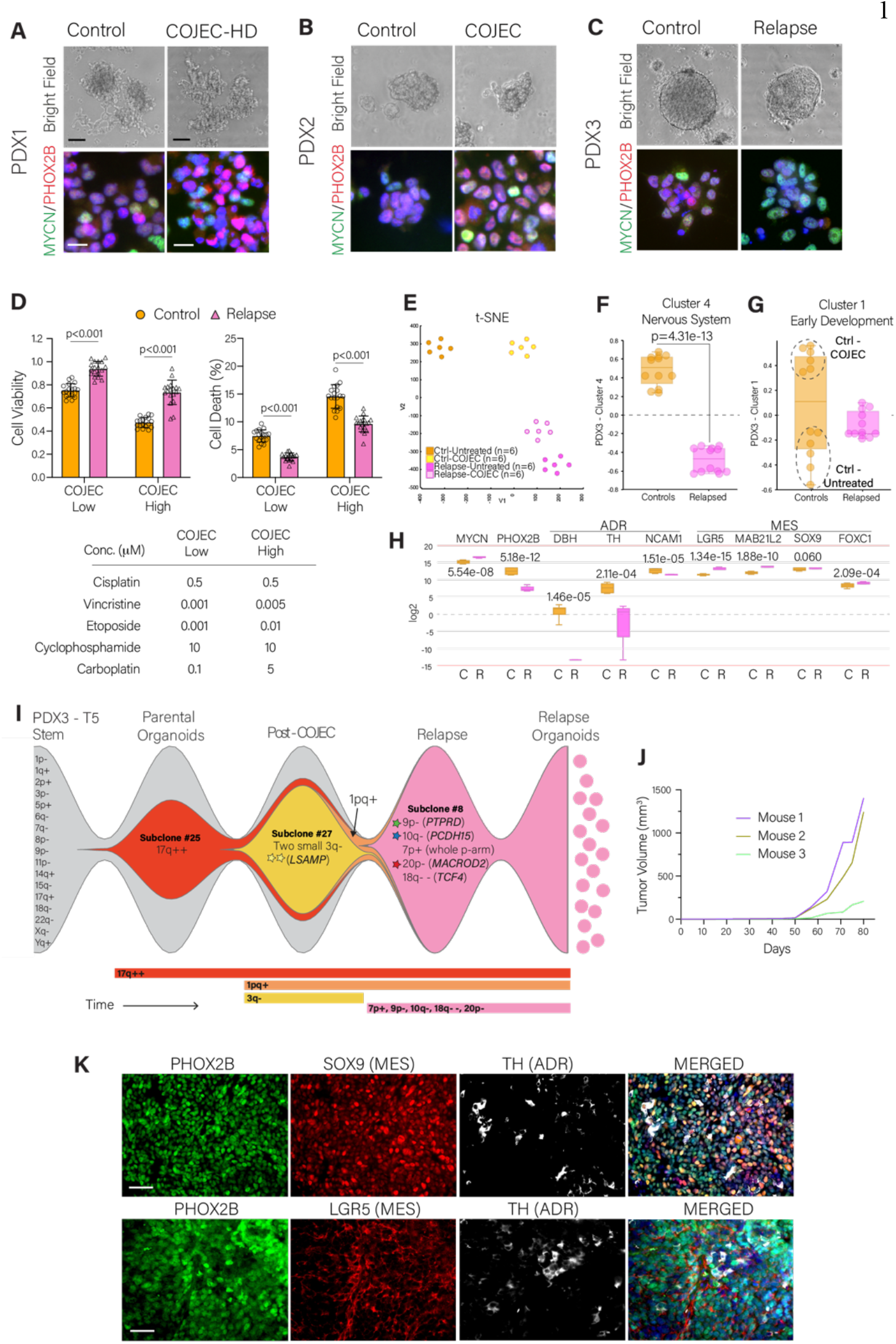
Establishment and characterization of COJEC-treated NB PDX-derived organoids. (**A-C**) NB organoids from control and COJEC-treated PDX tumors. Fluorescence staining of MYCN (green) and PHOX2B (red); scale bars, 100 μm (bright field) and 20 μm (fluorescence). (**D**) Cell viability and cell death (%) for PDX3-derived organoids treated with low or high dose COJEC *in vitro.* (**E**) Transcriptomic t-SNE plot of PDX3-derived organoids treated or untreated *in vitro* with COJEC High (as defined in D). (**F, G**) Expression of PDX3-Clusters 1 and 4 signatures in PDX3-derived organoids. Dotted line circles in (G) mark the untreated and COJEC-treated groups for control organoids. (**H**) Expression of selected genes in control (C) and relapse (R) PDX3 organoids. (**I**) Fish-plot depiction of the genetic evolution of a PDX3 COJEC-treated tumor, from the parental organoids, through the post-COJEC (surgical resection timepoint) and relapse stages, to the establishment of new organoids. Clones match those in Fig 2I-J. Subclone frequency is reflected by the relative size of colored region at specific timepoints. (**J**) Tumor growth of PDX3-relapse organoids in NSG mice (n=3). (**K**) Immunohistochemical fluorescence staining with PHOX2B (green), SOX9 (red, top panel), LGR5 (red, bottom panel), and TH (white). Nuclear staining with DAPI (blue). Scale bars, 50 μm.

We developed two ex vivo COJEC treatment protocols and applied these five chemotherapies combined to NB organoids derived from PDX3 tumors. Organoids derived from the relapsed T5 tumor displayed higher cell viability and lower cell death under COJEC treatment compared to control NB organoids (Fig. 7D). This suggested that NB organoids derived from COJEC-treated and relapsed NB PDXs can retain chemotherapy treatment resistance *ex vivo.*

We performed RNA-seq of the PDX3 control and relapsed organoids (untreated and COJEC-treated *ex vivo*) to examine if the organoids maintained transcriptional signatures similar to those observed *in vivo*. Each sample clustered well within their respective groups (Fig. 7E). We applied our previously identified PDX3 Cluster 4 (nervous system) and Cluster 1 (early development) gene signatures. Relapsed NB organoids displayed lower expression of the Cluster 4 signature compared to controls (Fig. 7F), consistent with our *in vivo* data (Fig. 4G, 5A). COJEC-treated control organoids showed significantly higher Cluster 1 expression than their untreated counterparts (Fig. 7G, Fig. S10D, p=1.94e-06).

We analyzed mRNA expression of key NB genes in all control organoids and relapsed organoids. *MYCN* expression was significantly higher in organoids derived from relapses compared to controls (Fig. 7H), consistent with the in vivo results. The expression of *PHOX2B* and the ADR markers *DBH, TH,* and *NCAM1* was lower in relapsed organoids compared to control organoids (Fig. 7H). The MES markers *LGR5, MAB21L2, SOX9,* and *FOXC1,* which we identified from the in vivo data, maintained an overall higher expression in relapsed organoids compared to controls (Fig. 7H). Nonetheless, within the control organoids, we observed significantly higher expression of some individual MES markers in the COJEC-treated group (Fig. S10E), consistent with the higher expression of Cluster 1 in this group (Fig. 7G, Fig. S10D).

DEG analysis showed that COJEC-treated control organoids presented with downregulation of cell cycle genes along with upregulation of genes in specific morphogenesis and cell motility pathways (Fig. S10F-S10G). For relapse organoids, the main DEGs identified between the treated and untreated groups were related to cell cycle (Fig. S10H-I).

Using the relapse organoid derived from PDX3 T5, we confirmed by scDNA-seq that genetic aberrations were maintained from in vivo to ex vivo (Fig. 7I). The relapsed PDX3 T5 was a clonal sweep of subclone #8 (Fig. 2I). The convergent smaller deletions of genes *PTPRD* and *MACROD2,* as well as the deletion of *PCDH15* (detected in subclone #8, as well as several PDX3 tumors after treatment), were present in the COJEC relapse organoids and consistently maintained in the established organoids. Thus, the selective sweep of subclone #8 in the relapse was complete and no new aberrations were detected in the organoids (Fig. 7I). The subclone with *LSAMP* deletion (#27, yellow) occurring in approximately 60% of the T5 tumor at the time of surgical resection, was not retained neither in the relapse nor the organoids, illustrating continuous clonal evolution over time.

Finally, we injected the relapsed NB organoids subcutaneously into NSG mice (n=3), and showed that they maintain tumorigenic capacity in vivo (Fig. 7J). Tumors displayed an undifferentiated morphology, were positive for the NB marker PHOX2B, and maintained high amounts of MES markers SOX9 and LGR5 along with low ADR marker TH (Fig. 7K), in concordance with the parental relapse tumor (Fig. 5D).

Thus, NB organoids derived from COJEC-treated relapsed PDXs exhibited the following properties: (i) display increased resistance to chemotherapy ex vivo; (ii) maintain transcriptional signatures of their corresponding in vivo tumors; (iii) are genetically similar to the in vivo tumor; (iv) are tumorigenic, and (v) maintain relative expression of ADR and MES-associated markers.

## DISCUSSION

Treatment resistance and relapse are common for patients with high-risk NB. Chemoresistance is usually studied *in vivo* by applying a single drug treatment (for example, cisplatin). Here, we developed a clinically relevant COJEC-like treatment protocol including the five chemotherapeutic agents usually given as first-line treatment to children with high-risk NB. This protocol was applied to multiple NB *MYCN*-amplified PDX models, and genomic and transcriptomic analyses were performed at different time points through the treatment course. Our analyses revealed substantial genetic diversity, as well as high intra- and inter-tumor heterogeneity, at both the DNA and transcriptional levels. NB PDXs in mice with upfront progression during chemotherapy and relapsed NB PDXs after chemotherapy showed an immature transcriptional phenotype that resembles the previously described MES-like cell state (*10, 12, 15–17, 19*). In contrast, NB tumors from mice that were cured after COJEC and surgical resection showed, at the time of surgical resection, features of nervous system development that resemble the ADR cell state (*10, 12, 15–17, 19*). The 3D NB organoids derived from relapsed PDXs retained chemo-resistance, transcriptional and genetic signatures of the in vivo tumor from the mice, as well as tumorigenicity, showing that these features can be maintained ex vivo.

Although a mutation-centric view has been dominant for years, recent data also point to transcriptionally-plastic oncogenic cell states as mediators of cancer development and treatment resistance (*42–44*). Several studies have demonstrated the existence of at least two transcriptional cell states in NB: an immature MES-like cell state and an ADR differentiated cell state (*10–13, 15–19*). There is consensus across studies for the ADR cell state, but inconsistencies exist in the definition of the immature MES-like state, with some data even questioning the existence of this cell state beyond cell culture (*12, 45*). The discordances observed between studies could be due to differences in methodology, including technical and bioinformatic methods, or sample material, such as cell lines, mouse models, PDXs, or patients, or the use of treatment naïve vs. treated tumors.

Our findings showed that these transcriptional cell states occur in patient-derived *MYCN*-amplified NB models both in vivo and ex vivo. Our results also demonstrated the relevance of the immature MES-like cell state in intrinsic and acquired NB treatment resistance. In vivo and ex vivo data suggested that COJEC treatment could select for cells with the immature MES-like phenotype and that these cells might contribute to relapse. In contrast, a predominantly ADR signature right after COJEC treatment correlated with the absence of relapses and a good overall prognosis. Our results also identified multiple cells positive for both ADR and MES markers (Fig. 5D and 7K), indicating the presence of cells in an intermediate state, which has been previously proposed (*10, 11, 17, 19*). In addition, we showed that the time point of transcriptional analysis is important; for responsive tumors, the immature phenotype was primarily identified in the relapses, and this phenotype was only detected in the absence of treatment in tumors with upfront progression. Our results are consistent with findings in which an NB cell subtype with immature mesenchymal characteristics was enriched in relapsed patient tumors (*10, 16*). Our data also suggested that transcriptional analysis and histochemical evaluation of ADR and MES markers, in addition to genomic assays, may be clinically valuable when assessing NB prognosis and treatment course. In particular, at the surgical resection post-COJEC timepoint, when specific signatures (cell cycle, ADR, MAPK) and markers (SOX9, LGR5) could provide valuable information about the COJEC response. Additionally, our results pointed to the importance of targeting both cell states from the start of treatment, to address upfront progression and prevent relapses.

We identified examples of convergent evolution between the PDX models, parallel evolution within each PDX model, clonal sweeps, and several examples of smaller deletions of genes that regulate nervous system development and chromosomal stability, both of which may influence chemoresistance. Convergent evolution of small deletions in two or more of the PDX models indicated that Darwinian selection occurred during tumor progression. However, these changes were detected in both controls and treated tumors, suggesting that they are important for NB progression but not necessarily for treatment resistance. We found no recurrent CNAs that could explain resistance or relapse. Thus, although the common NB CNAs lay the oncogenic foundation for NB growth, the development of relapses seems to be primarily mediated by transcriptional (and likely epigenetic (*16*)) changes. Our findings that COJEC-resistance was influenced by the immature MES-like phenotype while tumors lacked specific clonal selection at DNA level suggested that treatment-resistant relapses can develop as a result of Lamarckian induction, rather than through pure Darwinian selection. The level of (in)stability of the relapse-associated immature phenotype will be important to understand. The phenotype could, in principle, become stable through acquisition of genetic changes. However, we did not identify specific CNAs that could confer such stability. The identification of the immature phenotype in PDX tumors weeks after cessation of treatment in vivo and in organoids established ex vivo indicated a certain level of intrinsic stability of the phenotype even without treatment pressure.

A limitation in our study is that we analyzed a limited set of NB PDX models and, given the broad heterogeneity of NB, it remains to be shown if our findings represent a general phenomenon of all *MYCN*-amplified NBs or only a subset of tumors. We did not investigate *MYCN* non-amplified high-risk NB because of a lack of stable PDX models for this subtype. Our studies were performed using immunodeficient mice and the immune system can interact with the NB cell phenotypes (*46, 47*), which could yield a different outcome in fully immunocompetent individuals. On the other hand, our findings demonstrated that NB can adopt an immature MES-like phenotype even in the absence of a complete immune system.

The in vivo confirmation of two NB phenotypic cell states opens for novel therapeutic opportunities because current treatment protocols are not designed to target different NB cell states. It is conceivable that the cell states are not binary, but NB cells exist along a continuum of states. Our results showed the relevance of the MAPK and ERK cascades, in line with findings that suggest therapeutic targeting of these pathways in chemotherapy-resistant NB (*16, 48, 49*). Additionally, other work shows that Tumor Necrosis Factor Related Apoptosis-Inducing Ligand (TRAIL) therapy (*21*) and immunotherapy (*46*) could also be of interest. Given the clinical heterogeneity in NB patient tumors, and reflected in our NB PDX treatment responses, treatment resistance can likely develop by multiple mechanisms in individual tumors, as demonstrated in melanoma (*7*). Nevertheless, successful treatment of NB will likely need to target both, or multiple, cell states. Future studies need to define how and when specific targeting of the NB cell states should be implemented.

## MATERIALS AND METHODS

### Experimental Design

The goal of this work was to uncover the mechanisms involved in COJEC treatment resistance in high-risk NB. We designed and tested a COJEC-like protocol using three NB PDX models. In vivo, mice were injected subcutaneously with dissociated PDX organoids and, upon tumor engraftment, the mice were randomly allocated to treatment groups aiming for a minimum of 5 mice per group (on the basis of previous experimental experience). In vivo experiments were not blinded. Mice were euthanized based on tumor size, weight loss, overall health deterioration, or end of study time. No tumor-bearing mice were excluded. Only one outlier was identified (PDX3-T10), which was considered as part of the COJEC (only) group for mice survival analysis and as part of the PC group (not cured) for subsequent tumor analyses. Samples were blinded for morphological differentiation assessment. Bulk DNA and RNA were collected from the same piece of tumor tissue to allow for direct comparison. SNP and scDNA-seq were performed for samples of interest based on tumor response. RNA-seq was performed for all tumors. All RNA analyses were performed using the embedded analysis tools within the R2 Genomics Analysis Platform and Metascape. For in vitro experiments, three biological replicates with three technical replicates each were performed. For the organoids RNA analysis, two biological replicates with three technical replicates each were analyzed.

### Animal Experiments

All procedures were conducted according to the guidelines from the regional Ethics Committee for Animal Research (permit no. M11-15 and 19012-19). NMRI nude mice were purchased from Taconic and NSG mice were obtained from in-house breeding. NB PDX dissociated organoids (2×10^6^ cells) were suspended in a 100 μl mixture (2:1) of stem cell medium and Matrigel (Corning, Cat No.354234) and injected subcutaneously into the flanks of female mice. Mouse breed was selected based on the tumor engraftment rate for each PDX; PDX1 was analyzed in both nude (n=9) and NSG mice (n=24), PDX2 in NSG mice (n=22) and PDX3 in nude mice (n=41). Tumor size was measured using a digital caliper and calculated with the formula V=(πls^2^)/6 mm^3^ (l=long side and s=short side of each tumor). Mice were randomly allocated to control, cisplatin or COJEC groups once their tumor had reached approximately 500 mm^3^. The cisplatin group was treated intraperitoneally with 4 mg/kg cisplatin dissolved in sterile saline (Selleckchem, Cat No.S1166) each Monday, Wednesday and Friday, the control group was treated with equal volume of saline. The COJEC group was treated with intraperitoneal injections of cisplatin (1 mg/kg; Selleckchem, Cat No.S1166) and vincristine (0.25 mg/kg; Santa Cruz, sc-201434) on Mondays, etoposide (4 mg/kg; Santa Cruz, sc-3512) and cyclophosphamide (75 mg/kg; Santa Cruz, sc-361165) on Wednesdays, and carboplatin (25 mg/kg; Santa Cruz, sc-202093A) on Fridays. A subset of PDX1 mice (n=7) was treated with a high-dose COJEC protocol: cisplatin 1.5 mg/kg, vincristine 0.5 mg/kg, etoposide 6 mg/kg, cyclophosphamide 100 mg/kg, and carboplatin 35 mg/kg. Treatment was administered for a maximum of six weeks, making a maximum of 18 injection days per mouse. Treatment response was defined by the parameters established by the Pediatric Preclinical Testing Program (*50*) presented in Table S1. All mice were monitored for weight loss and other signs of toxicity. Treatment was paused if weight loss >10% to allow for recovery. When the tumor size had decreased to approximately 200 mm^3^, a subgroup of COJEC-treated PDX3 tumors (n=9) was surgically removed, treatment was interrupted and mice were monitored for tumor relapses. Mice were sacrificed when tumors reached 1800 mm^3^, or at a humane endpoint if weight loss >20% from initial weight or significant health deterioration was observed, or one year after initial cell injection. Upon collection, tumors were divided into pieces and stored for the corresponding experimental purposes.

### Immunohistochemistry and histological analyses

Tumors were fixed in 4% formalin, embedded in paraffin and cut in 4 μm sections for staining. H&E (Histolab Products) staining was performed for histopathological analysis. The slides were blinded for morphological differentiation assessment, and evaluated by an independent viewer. TUNEL assay (Abcam, ab206386), using either methyl green or hematoxylin as counterstain, was performed to asses cell death. Quantification was performed using the software QuPath 0.2.3. DAB staining was performed using the Autostainer Plus (Dako) and fluorescence staining was performed manually. Heat-induced antigen retrieval was done using sodium citrate buffer (pH 6.0). The antibodies used are PHOX2B (1:1000, Abcam), PHOX2B-AF647 conjugate (1:100, Santa Cruz Biotechnology), KI67 (1:200, Dako), CD34 (1:200, Santa Cruz Biotechnology), MYCN (1:200, Proteintech), TH (1:1000, Abcam), SOX9 (1:500, Abcam) and LGR5 (1:150, Abcam). Processing and quantification of fluorescence images was performed with Image J.

### Bulk DNA and RNA extraction

Snap-frozen tumors were added to RNAlaterTM-ICE (Invitrogen, #AM7030) and kept overnight at −20°C before extraction. DNA and RNA were extracted from tumors and organoids using the AllPrep DNA/RNA MiniKit (Qiagen, #80204).

### SNP-array DNA genotyping and analysis

DNA from selected PDX-samples was subjected to the Cytoscan HD array (Affymetrix) platform according to standard methods. The obtained CYCSPH-files were imported into Chromosome Analysis Suite (ChAS, Affymetrix, v.4.1.0.90 r29400). All samples from the same PDX were imported simultaneously and the chromosomes were assessed manually. Only genetic alterations clearly visible by eye, ≥ 50 kBp and with a marker count ≥ 50, were included in further analysis. Constitutional copy number variants (CNVs) were omitted based on manual comparison against the Database of Genomic Variants (DGV) with genome GRCh37/hg19. For each genetic alteration, its genomic location, type of aberration (loss/gain) and logarithmic median probe intensity ratio (log_2_ R) was extracted. The function Rawcopy (v.1.1) (*51*) in R was used on the Affymetrix fluorescence array intensity (CEL) files and its output was used to generate Tumor Aberration Prediction Suite (TAPS, v.2.0) (*52*) plots of copy number clusters for each chromosome and sample.

Each detected genetic alteration was analyzed in unison with the TAPS-plots to determine the allelic composition and the clonality (clonal/subclonal) for the aberrations. The log2 R together with the ploidy of the genetic alteration (N_t_) as well as the background cells (N_p_), were used to compute the mutated sample fraction (MSF), defined as the percentage of cells in that sample that harbor this alteration.

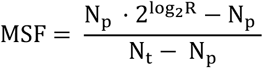

For cnni:s, the mirrored B-allele frequency (mBAF) together with information about its allelic composition (number of A- and B-alleles, N_A_ and N_B_) was used for MSF computation.

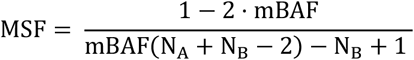

The MSF was used to compute the mutated clone fraction (MCF) defined as the proportion of cells in the sample having a specific genetic alteration divided by the purity or tumor cell fraction (TCF) of that sample. The TCF was computed for each sample by calculating the mean of the MSF-values of all aberrations in a sample defined to be clonal by the visual inspection of TAPS-plots. The interval of clonal events was computed using the standard deviation of the MSF-values of the alterations used for TCF-computation (SD_MSF_) as

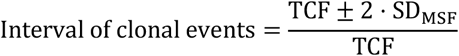

defined as the MCF-interval in which an alteration is determined as clonal. If a genetic alteration has a MCF in that sample that is within the interval of clonal events, it was set to 100 %, otherwise the MCF-value were kept, and the event was defined as subclonal.

### Single-cell low pass CNA sequencing

Single cells were prepared from viably frozen tumors by mechanical dissociation with sterile scalpels and enzymatic dissociation with Liberase DH (0.15 mg/mL) at 37°C and 250 rpm for 20-40 min depending on tumor size. Then, tumor cells were treated with DNase I (2.5 μg/ml, Sigma-Aldrich), trypsinized (0.05%) and filtered through a 70 μm cell strainer. Single-cell whole genome sequencing (scDNA seq) was performed as previously described (*53–55*). Library preparation was performed from each isolated nucleus. Low-pass (0.01-0.02x) whole genome sequencing produced reads which were aligned to human reference genome (GRCh38). Copy numbers were determined by AneuFinder (version 1.8.0) at 1 Mb bin sizes. Analysis quality was determined as previously described (*53*). Manual curation of single cell CNA profiles identified true imbalances based on the previously optimized cut-off of five consecutive 1Mb bins (*33*). High-grade amplifications of *MYCN* (2p24.3-24.2) and of 4q28.3-q31.1 were scored based on two consecutive 1Mb bins. Specific clones based on unique CNAs were identified for all single cells within the respective tumor. In total, 18 samples (19-85 libraries (cells)/tumor) were successfully analyzed. This resulted in a matrix where each row is a bin, approximately 1 Mbp of size, each column is a single cell, and the matrix elements are the number of copies of that particular segment. The matrix was used as input to an algorithm written in R. This information was used to construct an event matrix where each row is a genomic event, and each column is a cell or group of identical cells.

### Phylogenetic and genomic analyses

To analyze the evolutionary trajectory of the genomic alterations across the PDX-samples, DEVOLUTION (*56*) was used. The input is a matrix in which each row represents a genetic alteration in a particular sample, along with information about that genetic alteration’s genomic location, type of alteration and MCF. Hence, for bulk genotyping data we obtain an event matrix using DEVOLUTION and for single cell whole genotyping data we obtain an event matrix using a customized algorithm written in R. The event matrices obtained from DEVOLUTION for bulk data and from the customized algorithm for single cell data were used to reconstruct phylogenetic trees using the maximum likelihood and maximum parsimony methods using the phangorn (v2.8.1) (*57*) package and visualized using the ggplot2 (v.3.3.5) package.

Genetic diversity was calculated using multiple metrics based on clonal/subclonal information and phylogenetic analysis. First, the fraction of private aberrations (clone/subclone detected in only one tumor within the PDX model) gave a measure of inter-tumor heterogeneity. Second, copy number aberration burden (CNAB) was calculated from the number of accumulated copy number aberrations (CNAs), to describe the total burden of CNAs within each tumor:

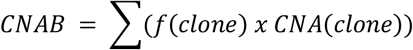

where f(clone) is the frequency of each clone/subclone in one tumor and CNA (clone) is the absolute value of the copy number deviation compared to a diploid cell. The sum is over all genetic alterations and all subclones encompassed by that tumor. To allow for comparison between treatment groups within each PDX, only CNAs that were not present within the stem of that PDX were included in the calculations and graphs. Third, genetic diversity was calculated with inverse Simpson’s index of diversity (Ds) as:

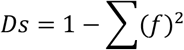

where f equals the frequency of each clone. To visually depict the frequency of clones at certain time points and the emergence of new clones over time, a fish plot based on CNA events at specific time points during treatment was produced using an R script (*58*).

Comparison between groups and test for trend for the genetic diversity measures was done by one-way ANOVA. The correlation between CNAB and tumor collection day was investigated with Spearman correlation and linear regression. All statistical analysis was performed by the GraphPad software.

### RNA sequencing and analysis

mRNA library preparation was performed using Illumina Stranded mRNA Prep, Ligation (Illumina, #20040534) and TruSeq® Stranded mRNA Library Prep (Illumina, #20020594) kits on the King Fisher FLEX system (Thermo Fisher Scientific, #18-5400620). Sequencing was done using the NovaSeq 6000 SP Reagent Kit v1.5 (200 cycles, Illumina, #20040719) and the NovaSeq 6000 S1 Reagent Kit v1.5 (200 cycles, Illumina, #20028318) on the NovaSeq 6000 System (Illumina, #20012850). Reads were aligned to the Human GRCh38 reference genome (Ensembl) using STAR (*59*) and transcript summarization was carried out using featureCounts with subread package (*60, 61*) using the annotation (GTF) from Gencode version 33. Expression count matrixes from the different sequencing runs were merged and batch corrected using CombatSeq (*62*), followed by variance stable transformation (VST) of the resulting matrix using DESeq2 R package (*63*). Library preparation, mRNA sequencing, data normalization and batch correction were performed by the Center for Translational Genomic, Lund University. The data is available on the R2: Genomic Analysis and Visualization Platform (http://r2.amc.nl) under__(upon acceptance for publication).

All RNA-seq analyses were performed on the R2 Genomics Analysis and Visualization Platform. The Kocak dataset, was used for patient overall survival analysis (Stages 3-4, n=219, median cut). Visualization of t-SNE plots, heatmaps and volcano plots was done using R2, FDR<0.01. Box plots for genetic signatures present the average z-score values over the gene set for each sample. Welch’s t-test correction was used to analyze statistical differences between groups for individual signatures. Gene ontology (GO) enrichment analysis was performed using the Gene Ontology Platform (*64–66*) and Metascape (*67*). Visualization of GO enrichment and DEG was performed using Revigo (*68*) and Metascape.

### *In vitro* cultures

NB PDX organoids previously established derived from PDX1, 2 and 3 (*22, 23, 69*) and newly established organoids, were cultured as free-floating tumor organoids under serum-free conditions in Dulbecco’s modified Eagle’s medium and GlutaMAX F-12 in a 3:1 ratio, supplemented with 1% penicillin/streptomycin, 2% B27 without vitamin A, fibroblast growth factor (40 ng/ml), and epidermal growth factor (20 ng/ml). Cells were tested for Mycoplasma prior to experiments. The identity of the NB PDX cells was confirmed using STR/SNP analysis.

New PDX-derived tumor organoids were established by performing mechanical dissociation of fresh pieces of tumor using a sterile scalpel, followed 24 hours later by enzymatic dissociation with Liberase DH (0.15 mg/ml, Roche Ref. 05401054001).

### Cell viability and cell death assays

Cell viability and death were measured using the CytoTox-Glo Cytotoxicity Assay (Promega). Organoids were dissociated using Accutase (Sigma) and single cells were seeded and treated in triplicates in white 96-well plates with clear bottom for 48 hours, for three biological replicates. Cells were incubated for 48 hours before treatment to allow the formation of small organoids. Luminescence readouts were performed using a Synergy2 plate reader (BioTek). Drugs used for *in vitro* treatment were the same used for the *in vivo* work.

### Immunofluorescence

Dissociated organoids were seeded on glass slides pre-coated with LN521-05 laminin (10 ug/ml, Biolamina). Cells were allowed to attach and grow for at least 72 hours, then they were fixed using 4% paraformaldehyde. Permeabilization and blocking was performed with 0.25% Triton X and 2.5% Fetal Bovine Serum in 1XTBS. The primary antibodies used were against MYCN (1:100, Santa Cruz Biotechnologies, sc-142) and PHOX2B (1:100, Abcam, ab183741). The secondary antibodies used were AlexaFluor 488 (1:200, Invitrogen, Ref. A11001) and AlexaFluor 633 (1:200, Invitrogen, Ref. A21071). Nuclear staining was performed with DAPI (Invitrogen, Ref. D3571).

### Statistical analyses

All statistical analyses pertinent to DNA and RNA analysis were performed as described in the corresponding methods sections. The remaining analyses were performed using GraphPad Prism 9. The log-rank test was used to determine significance for Kaplan-Meier survival curves for *in vivo* studies. Non parametric Mann-Whitney test was used for the morphological differentiation analysis, and samples were graded as undifferentiated (0), mixed (1) or differentiated (2). For the analysis of TUNEL cell death quantification, ordinary one-way ANOVA was performed with Dunnett’s test correction for multiple comparison by treatment group compared to controls. For tumor volume comparison, ordinary one-way ANOVA was performed with Welch’s t-test correction for multiple comparison. For the organoid drug testing, two-way ANOVA with Sidak correction for multiple comparison by concentration was performed. Overall, a p-value<0.05 was considered significant.

## Supporting information

Supplementary Figures - Manas et al

## SUPPLEMENTARY MATERIALS

Fig. S1. A COJEC-like protocol for the treatment of NB PDX models in vivo.

Fig. S2. Clonal dynamics and genetic diversity across NB PDX models.

Fig. S3. Clonal dynamics from NB PDX3.

Fig. S4. Phylogenetic analysis of low pass single cell (sc)DNA profiles across NB PDX models.

Fig. S5. Baseline transcriptional signatures for each NB PDX model.

Fig. S6. Characterization of transcriptional signature clusters for each NB PDX.

Fig. S7. DEG analysis and key transcriptional signatures across NB PDX3 treatment groups.

Fig. S8. Publicly available ADR and MES-like signatures across NB PDX3 treatment groups.

Fig. S9. Identification of phenotypic markers independent of stroma content and correlation of genetic aberrations and prognosis.

Fig. S10. Treatment and characterization of PDX-derived NB organoids.

Table S1. Individual responses to COJEC and Cisplatin treatment.

Data file S1. Complete lists of transcriptional signatures, DEG analysis and gene ontologies

Data file S2. Complete data for the Metascape enrichment analysis and network plots

## Acknowledgments

We thank the Center for Translational Genomics (CTG), Lund University and Clinical Genomics Lund, SciLifeLab for providing sequencing services, especially David Lindgren for his unvaluable assistance. We thank Jan Koster for his insightful comments. We thank Nancy R. Gough (BioSerendipity, LLC) for editorial services.

## Author contributions

AM, KA and DB conceptualized and designed the research;

AM, KH, AA, AS, HVB, KR and JE performed experiments;

AM, KA, NA, HY, MSB, DS, JK, DG and DB analyzed data;

FF, DG and DB supervised the research;

AM, KA and DB wrote the manuscript;

All authors reviewed the manuscript;

DB acquired funding and administered the project.

## Data and materials availability

RNA-seq data has been deposited to the R2 Genomic Analysis and Visualization Platform (http://r2.amc.nl) under the name:.........(upon acceptance for publication). The DNA-seq data has been deposited to.........(upon acceptance for publication). The raw code for the single cell analysis algorithm has been deposited at.....(upon acceptance for publication). All remaining data are available in the main text or the supplementary materials.

