## Supplementary Figures - Manas et al for "Clinically-relevant treatment of PDX models reveals patterns of neuroblastoma chemoresistance"

**One Sentence Summary:** COJEC chemotherapy treatment of neuroblastoma PDX models uncovers patterns of transcriptional plasticity and chemoresistance.

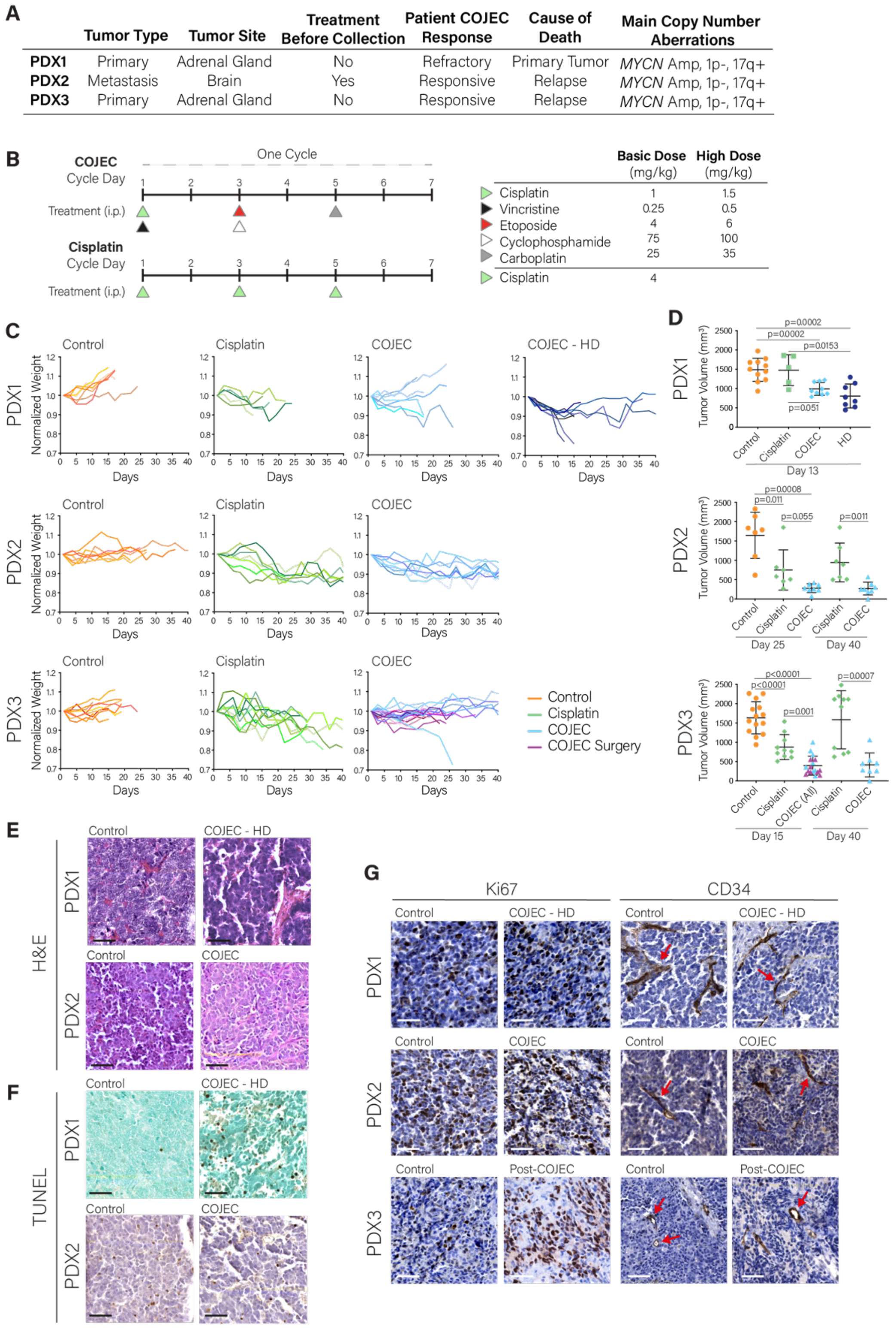


**Fig. S1. A COJEC-like protocol for the treatment of NB PDX models *in vivo*.** (**A**) Summary of general characteristics of the NB PDX models. (**B**) Dosing schedule for the COJEC-like cycle and the cisplatin cycle, and drugs doses for basic and high dose tiers. i.p., intraperitoneal. (**C**) Normalized mouse weight during treatment. Mice received a maximum of 6 cycles of treatment. (**D**) Tumor volume comparison across treatment groups at the relevant survival days (PDX1, day 13; PDX2, day 25; and PDX3, day 15) and at the end of treatment (day 40). The last available point was considered when mice had been euthanized before the day of comparison. Significance was determined by ordinary one-way ANOVA with Welch's t-test correction for multiple comparison. (**E**) Hoechst and eosin (H&E) staining of PDX1 and PDX2 to assess morphological differentiation. (**F**) TUNEL staining of PDX1 and PDX2 to assess cell death. (**G**) Ki67 (proliferation marker) and CD34 (blood vessel marker) immunohistochemical staining for each PDX model. Red arrows indicate representative blood vessels. Scale bars, 50 µm.

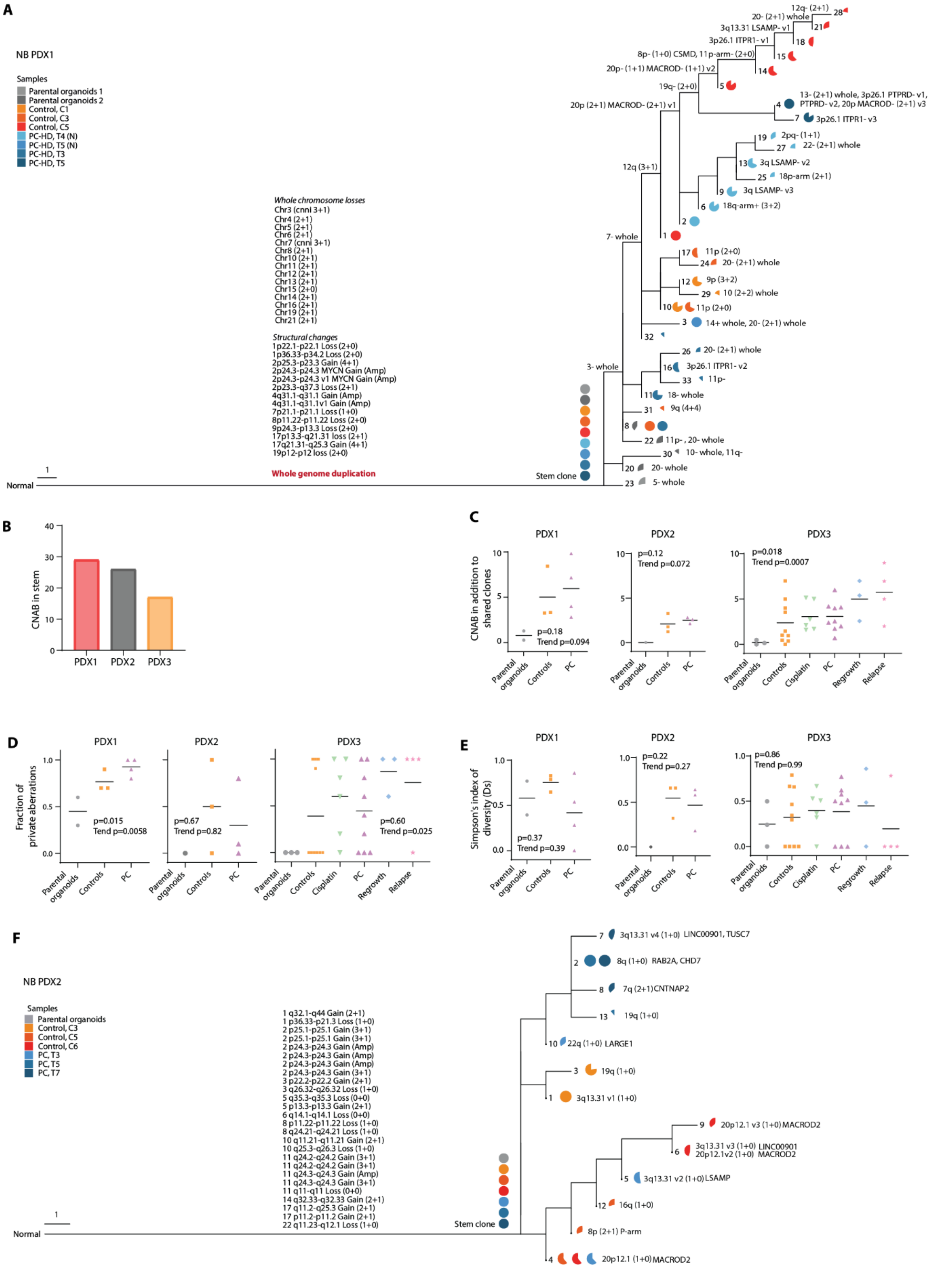


**Fig. S2. Clonal dynamics and genetic diversity across NB PDX models.** (**A**) Maximum-likelihood (ML) phylogenetic tree of NB-PDX1 based on CNAs. Frequency of different subclones is visualized with circle diagrams. Whole-genome duplication (WGD) early in the stem of NB-PDX1 was followed by a high number of large CNA events where the majority involved loss of the fourth copy of a region. Multiple occasions of parallel evolution were detected in NB-PDX1. (**B**) Number of CNAs in phylogenetic stem of each PDX model. (**C**) CNAB, (**D**) the fraction of private aberrations and (**E**) genetic diversity (Ds) per treatment group in PDX1, 2, and 3. Comparison between groups and test for trend by one-way ANOVA. (**F**) ML phylogenetic tree of NB-PDX2. Structural changes in the stem clone are listed.

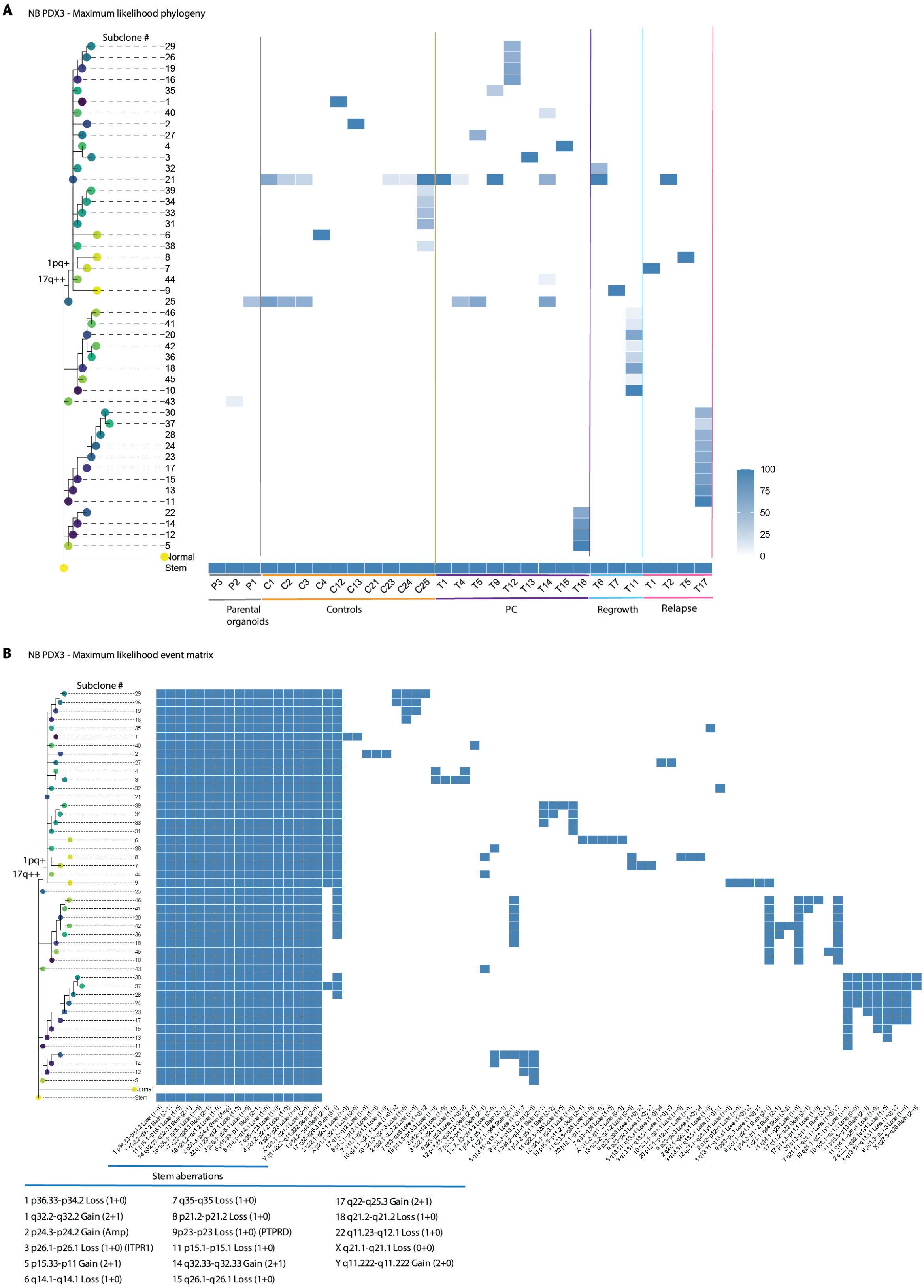


**Fig. S3. Clonal dynamics NB-PDX3.** CNA-based ML phylogenetic trees displaying all clones and subclones detected within PDX3. The trees in A and B are identical. The stem clone is defined as having all CNAs present in all organoids and tumors (lowest row in both trees). Subclones #1 – 46 are present in one or more tumors. The subclone number (#) corresponds to that in Fig. 2. Major branching events, 17q++ and 1pq+, are displayed in the trees. (**A**) The fraction of each subclone in the respective tumor is illustrated by the shade of blue. (**B**) Event matrix detailing the genomic location of the respective CNA in each subclone. The CNAs present in the stem are also written in larger text below the tree. The CNAs are presented as chromosome number, position range, type of aberration, copy number, and gene exhibiting convergent evolution, if applicable.

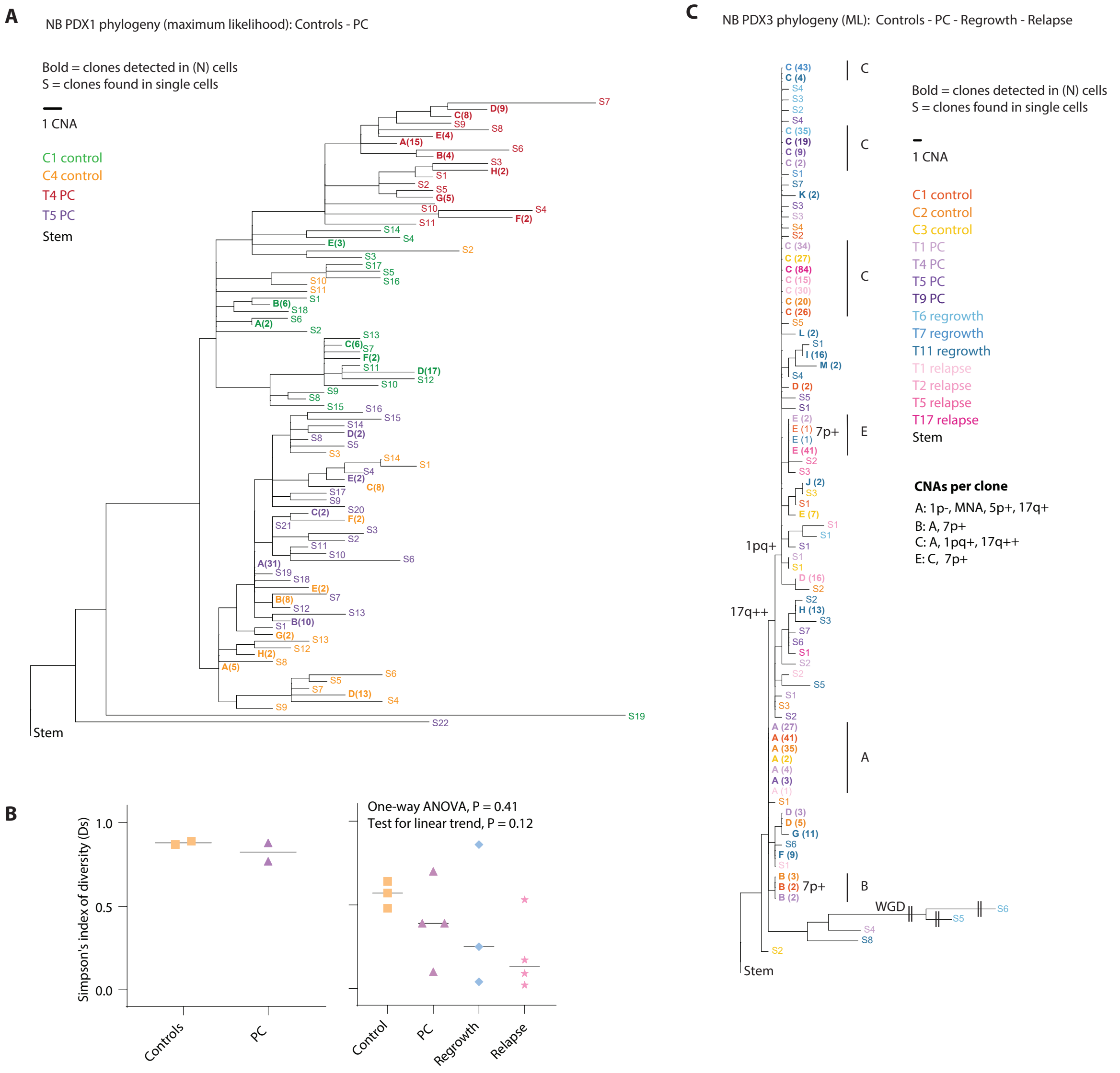


**Fig. S4. Phylogenetic analysis of low pass, single-cell (sc) DNA profiles across NB PDX models.** (**A**) ML phylogenetic analysis of NB-PDX1 scDNA. Clones labeled with "S" and a number represent single-cell clones that have a unique CNA profile from any other cell among the PDX tumor. Clones represented by multiple cells are indicated by bold letters with the number of cells detected indicated in parentheses. The clone text color reflects the PDX tumor (C1, C4, T4, T5) in which the clone was detected. All clones are unique to a single PDX1 tumor. (**B**) Genetic diversity (Ds) in tumors for the indicated treatment groups of PDX1 and PDX3. Comparison between groups and test for trend in PDX3 with one-way ANOVA. (**C**) ML phylogeny of NB-PDX3 scDNA. Major selective sweeps of 17q++ and 1pq+ are marked. In NB-PDX3, identical clones [A, B, C, E] were found in several of the tumors.

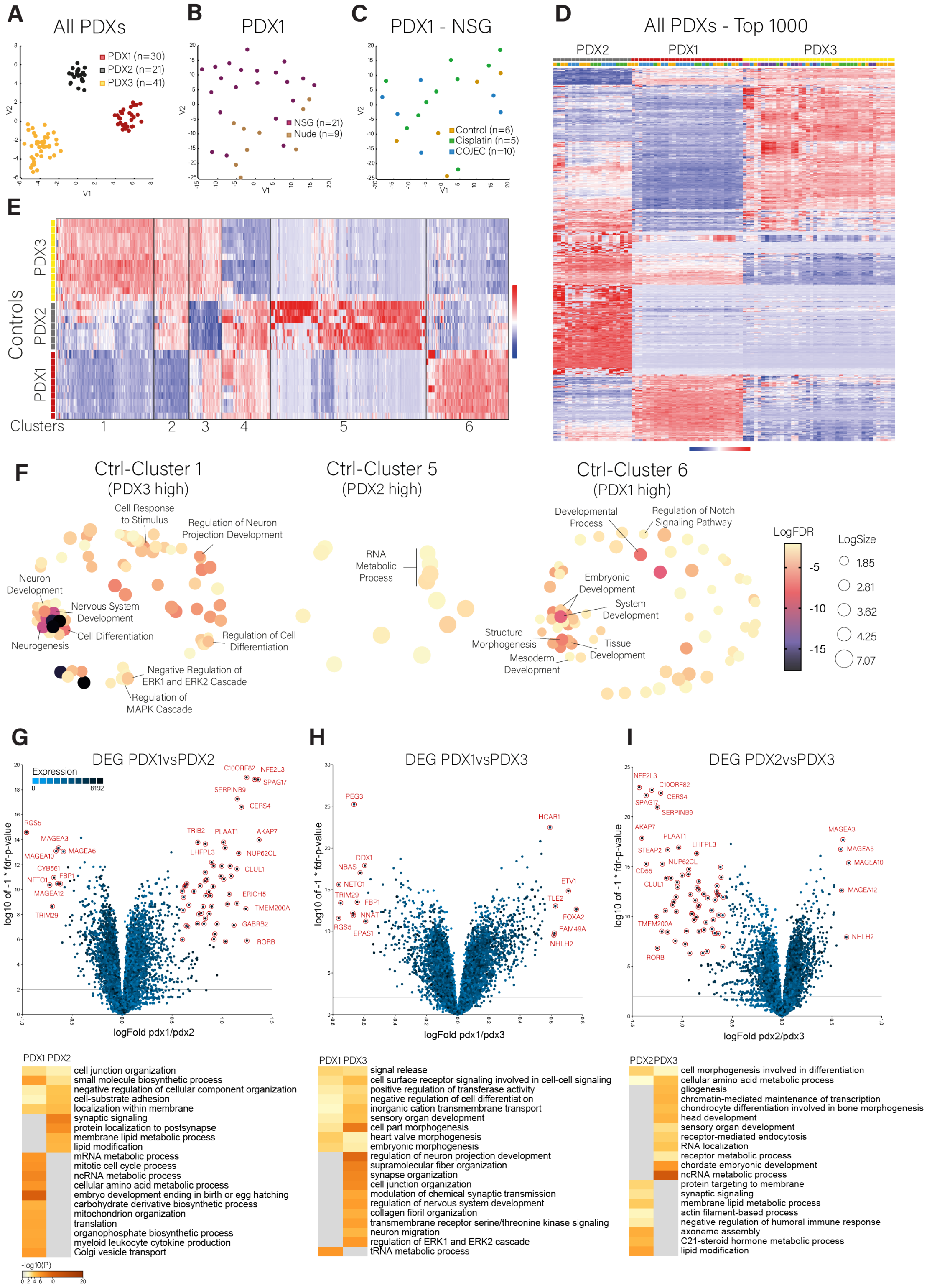


**Fig. S5. Baseline transcriptional signatures for each NB PDX model.** (**A-C**) t-SNE plots of all PDXs colored by PDX model (A), PDX1 colored by mouse strain (B), NSG mice PDX1 colored by treatment group (C). (**D**) Unsupervised top 1000 differentially expressed genes for all samples. (**E**) Unsupervised top 1000 differentially expressed genes for control samples. Six gene clusters are identified. (**F**) Multidimensional scaling (MDS) analysis and visualization of gene ontologies defined by Clusters 1, 5 and 6 from control (Ctrl) PDX samples in (E) (Revigo, non-redundant scaling 0.7). (**G-I**) Analysis of differentially expressed genes (DEG) between control samples of each PDX. Top panels display volcano plots of all DEG for each paired comparison (red genes = FDR < 0.01 and fold change > |1.5| , R2 Genomics Analysis Platform). Bottom panels display gene ontologies defined by the top 1000 DEG for each paired comparison (Metascape enrichment analysis for GO Biological processes).

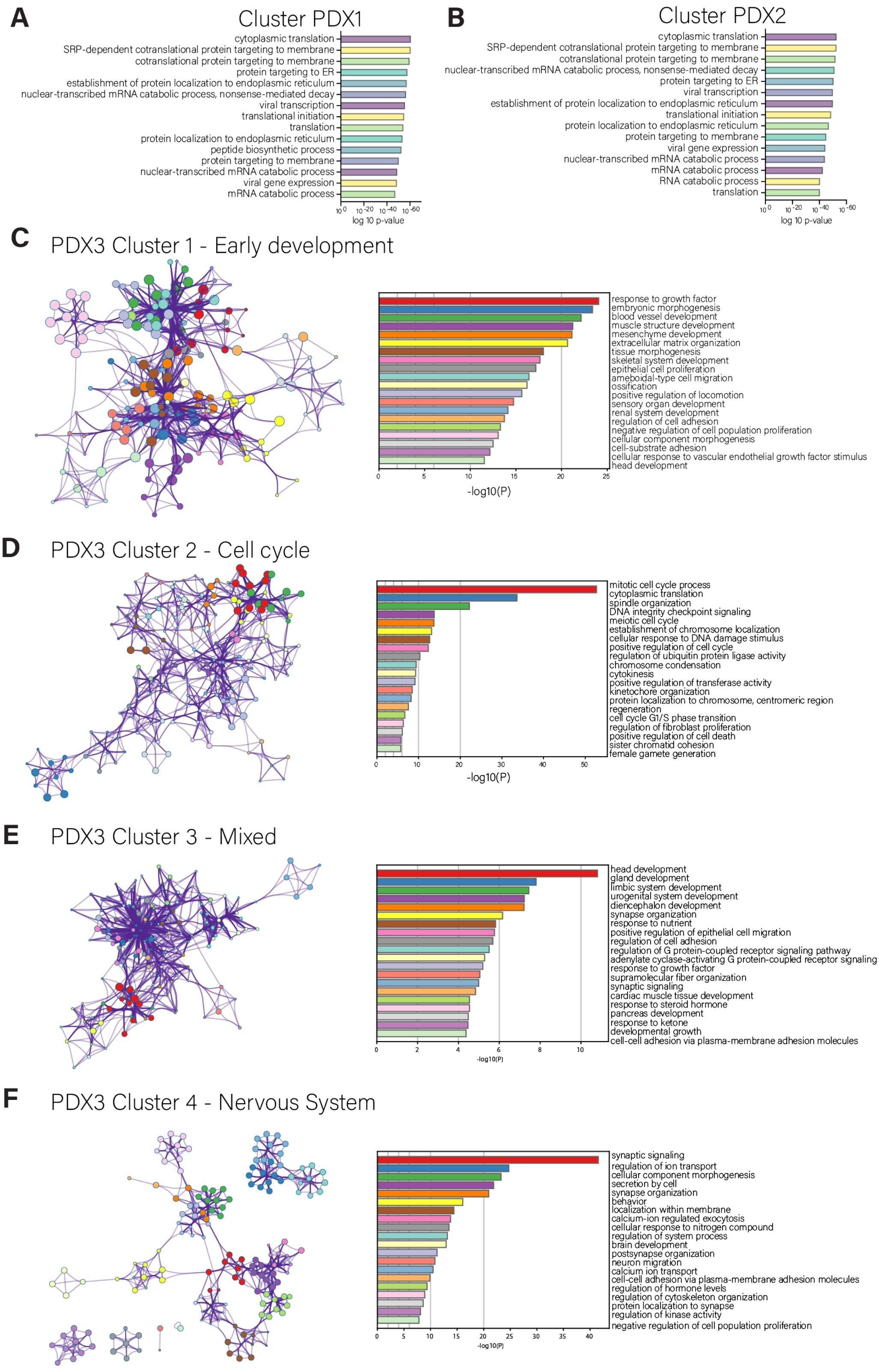


**Fig. S6. Characterization of transcriptional signature clusters for each NB PDX.** (**A-B**) Top 15 most significant gene ontologies defined by the single clusters identified for PDX1 and PDX2 (see Fig. 4E-F). (**C-F**) Enrichment analysis for the 4 clusters defined by PDX3 (Metascape enrichment analysis for GO Biological processes), visualized through network plots based on the top 20 most significant ontology clusters.

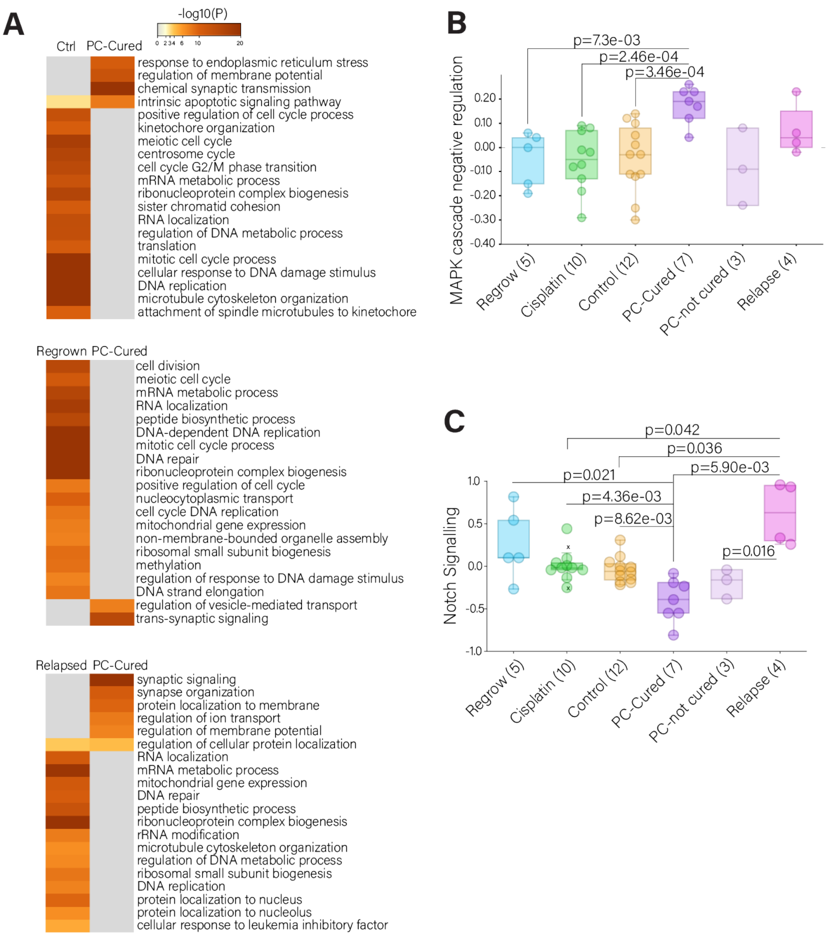


**Fig. S7. DEG analysis and key transcriptional signatures across NB PDX3 treatment groups**. (**A**) Gene ontologies defined by the top 1000 DEG between PDX3 treatment groups (Metascape enrichment analysis for GO Biological processes). (**B, C**) Statistical analysis with ordinary one-way ANOVA and Welch's t-test correction for multiple comparison of MAPK cascade negative regulation and Notch signaling across PDX3 treatment groups.

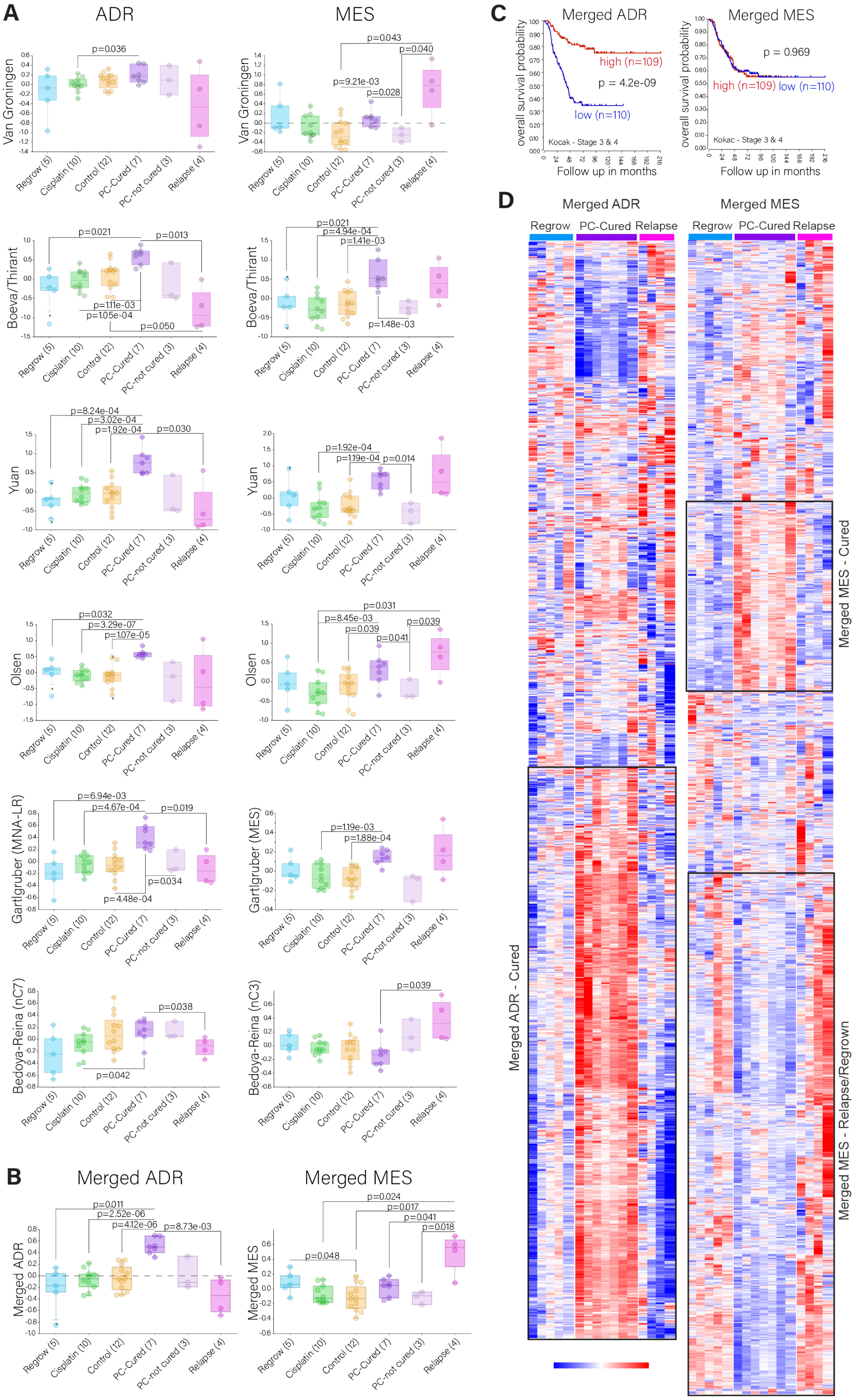


**Fig. S8. Publicly available ADR and MES-like gene signatures across NB PDX3 treatment groups.** (**A**) Analysis of six publicly available pairs of ADR-MES gene expression signatures *(10, 12, 15–17, 19)* across PDX3 treatment groups. (**B**) Analysis of the Merged ADR and Merged MES gene signatures (combining the 6 signature pairs from A, with PDX3 cluster 4 for ADR and cluster 1 for MES). (**C**) Overall survival Kaplan-Meier curves for the Merged ADR and MES gene signatures in high-risk patients (stages 3-4, Kocak dataset, R2 Genomics Analysis Platform). (**D**) Heatmap visualization of the Merged ADR and MES gene signatures, and cluster identification across the regrown, relapsed, and PC-cured sample groups.

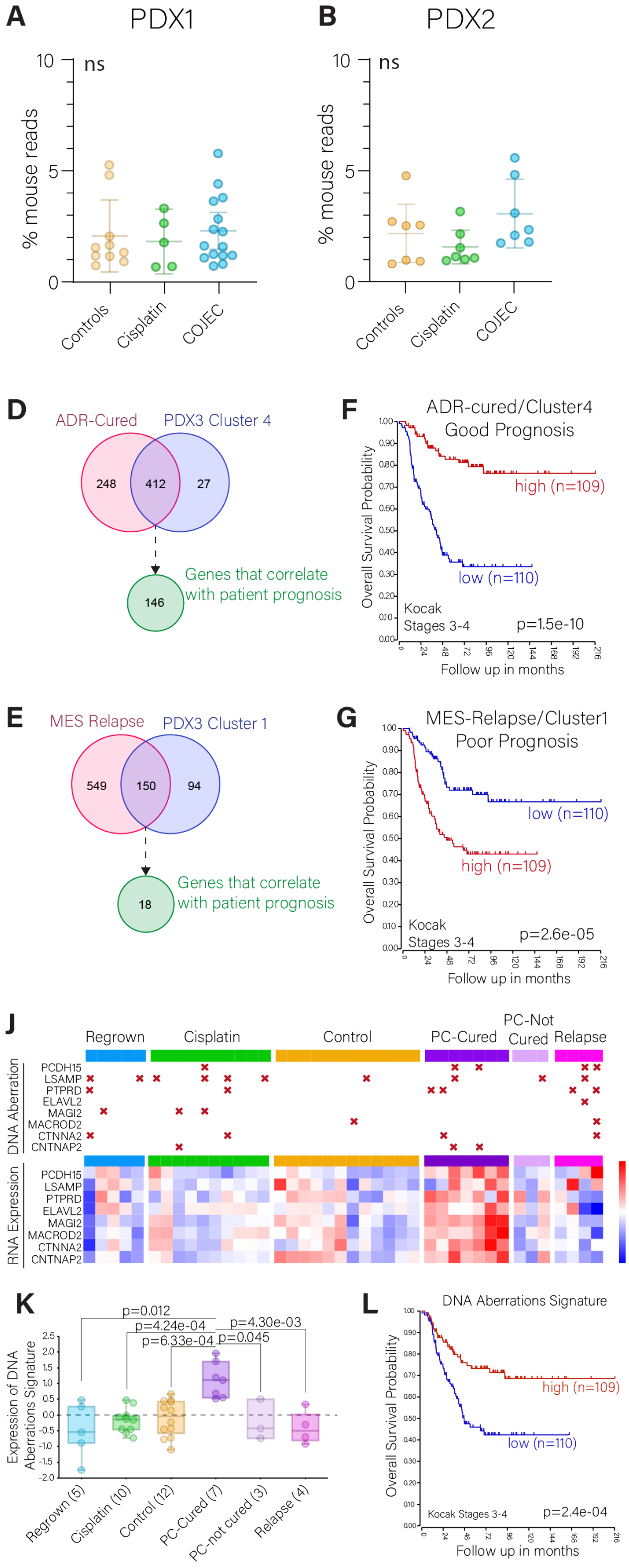


**Fig. S9. Identification of phenotypic markers independent of stroma content and correlation between genetic aberrations and prognosis.** (**A-C**) Percentage of mouse RNA reads in each tumor sample across NB PDXs. (**D, E**) Selection of ADR and MES genes common between the PDX3 defined clusters 1 and 4 and the Merged sub-signatures, and that individually correlate with patient prognosis (stages 3-4, Kocak dataset, overall survival, median cut, FDR < 0.05). (**F, G**) Overall survival Kaplan-Meier curves of the gene signatures from (D) and (E) on a cohort of 219 NB patients (stages 3-4, Kocak dataset). (**H, I**) Statistical analysis ordinary with one-way ANOVA and Welch's t-test correction of the gene signatures from (D) and (E) across the PDX3 treatment groups. (**J**) Comparison between convergent DNA aberrations (losses) and RNA expression of the corresponding genes for each PDX3 sample. (**K**) Statistical analysis of the signature from (J) across PDX3 treatment groups. (**L**) Overall survival Kaplan-Meier curves for the signature from (J).

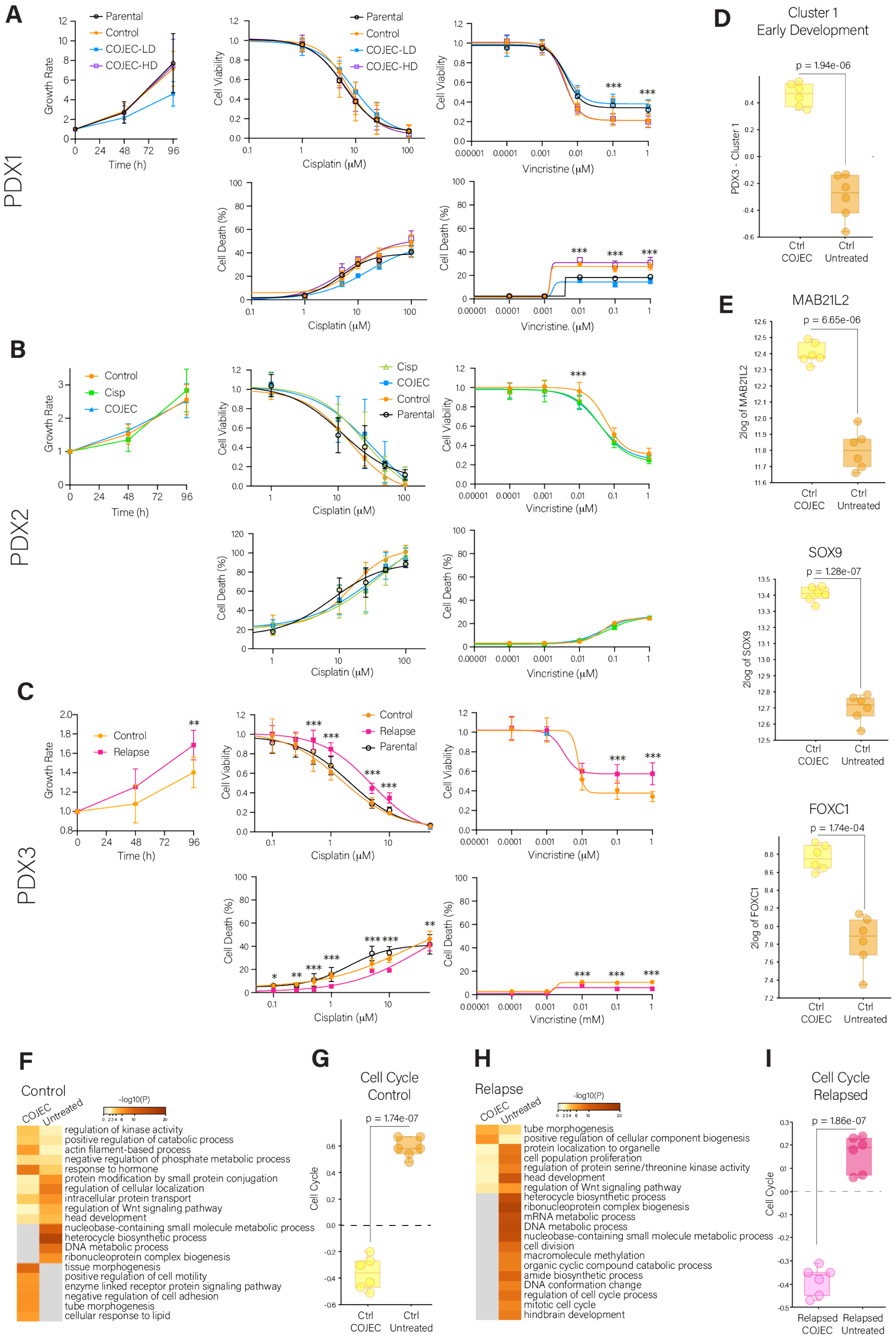


**Fig. S10. Treatment and characterization of PDX-derived NB organoids**. (**A-C**) Growth rate, cell viability, and cell death (%) for NB PDX-derived organoids treated with either Cisplatin or Vincristine as single drugs. Statistically significant differences were determined by two-way ANOVA with Sidak correction for multiple comparison (*, $p < 0.05$; **, $p < 0.01$, ***, $p < 0.001$). (**D, E**) Statistical analysis with Welch's t-test of Cluster 1 signature and expression of MES genes (*MAB21L2*, *SOX9*, *FOXC1*) with significantly different expression in COJEC-treated PDX3 control organoids compared to untreated organoids. (**F, H**) Gene ontologies defined by the top 1000 differentially expressed genes (DEG) between treatment groups for each set of NB organoids (Metascape enrichment analysis for GO Biological processes). (**G, I**) Statistical analysis with Welch's t-test of cell cycle gene expression between treatment groups for each set of NB organoids.

| Treatment Response | | |
|---|---|---|
| CR | Complete Response | Disappearance of tumor at least once |
| PR | Partial Response | Max Regression >50% |
| SD | Stable Disease | Max Regression <50%. Volume Increase at End of Treatment <25% |
| PD | Progressive Disease | Max Regression <50%. Volume Increase at End of Treatment >25% |
| - PD1 | Aggressive Progression | TGD<=1.50 |
| - PD2 | Delayed Progression | TGD>1.50 |
| TGD | Tumor growth delay | Survival day / Median survival day of controls |

| PDX1 | Mouse Strain | Max regression (%) | Vol. Increase at End of Treatment (%) | TGD | Response | Surgery | Relapse |
|---|---|---|---|---|---|---|---|
| T1 (N) | Nude | 0.00 | 209.37 | 1.60 | PD2 | No | |
| T2 (N) | Nude | 0.00 | 169.11 | 1.00 | PD1 | No | |
| T3 - HD (N) | Nude | 0.00 | 327.72 | 1.13 | PD1 | No | |
| T4 - HD (N) | Nude | 0.00 | 278.86 | 2.87 | PD2 | No | |
| T5 - HD (N) | Nude | 0.00 | 246.47 | 3.27 | PD2 | No | |
| T1 | NSG | 0.00 | 286.50 | 1.13 | PD1 | No | |
| T2 | NSG | 0.00 | 307.61 | 1.93 | PD2 | No | |
| T3 | NSG | 0.00 | 186.06 | 1.13 | PD1 | No | |
| T4 | NSG | 0.00 | 253.62 | 1.80 | PD2 | No | |
| T5 | NSG | 0.00 | 260.09 | 2.07 | PD2 | No | |
| T6 | NSG | 0.00 | 298.01 | 1.33 | PD1 | No | |
| T7 | NSG | 0.00 | 577.67 | 1.13 | PD1 | No | |
| T1 - HD | NSG | 0.00 | 168.44 | 1.00 | PD1 | No | |
| T2 - HD | NSG | 0.00 | 118.99 | 0.87 | PD1 | No | |
| T3 - HD | NSG | 0.00 | 315.89 | 2.40 | PD2 | No | |
| T4 - HD | NSG | 0.00 | 289.06 | 1.80 | PD2 | No | |
| T5 - HD | NSG | 37.81 | 19.75 | | SD | No | |
| Cis 1 | NSG | 0.00 | 240.43 | 1.00 | PD1 | No | |
| Cis 2 | NSG | 0.00 | 301.11 | 1.13 | PD1 | No | |
| Cis 3 | NSG | 0.00 | 383.02 | 1.60 | PD2 | No | |
| Cis 4 | NSG | 0.00 | 222.58 | 1.47 | PD1 | No | |
| Cis 5 | NSG | 0.00 | 226.16 | 0.87 | PD1 | No | |

| PDX2 | Mouse Strain | Max regression (%) | Vol. Increase at End of Treatment (%) | TGD | Response | Surgery | Relapse |
|---|---|---|---|---|---|---|---|
| T1 | NSG | 81.51 | -59.07 | | PR | No | |
| T2 | NSG | 47.31 | -8.16 | | SD | No | |
| T3 | NSG | 64.00 | -64.00 | | PR | No | |
| T4 | NSG | 45.78 | -24.94 | | SD | No | |
| T5 | NSG | 64.61 | -40.14 | | PR | No | |
| T6 | NSG | 73.27 | -53.18 | | PR | No | |
| T7 | NSG | 71.52 | -71.52 | | PR | No | |
| T8 | NSG | 100.00 | -100.00 | | CR | No | Yes |
| Cis 1 | NSG | 5.51 | 144.60 | 1.85 | PD2 | No | |
| Cis 2 | NSG | 57.70 | -19.26 | | PR | No | |
| Cis 3 | NSG | 1.10 | 60.63 | 2.04 | PD2 | No | |
| Cis 4 | NSG | 14.26 | -4.92 | | SD | No | |
| Cis 5 | NSG | 37.61 | 10.16 | | SD | No | |
| Cis 6 | NSG | 0.00 | 166.94 | 0.96 | PD1 | No | |
| Cis 7 | NSG | 30.36 | 35.96 | 2.31 | PD2 | No | |

| PDX3 | Mouse Strain | Max regression (%) | Vol. Increase at End of Treatment (%) | TGD | Response | Surgery | Relapse |
|---|---|---|---|---|---|---|---|
| T1 | Nude | 75.63 | -75.63 | | PR | Yes | Yes |
| T2 | Nude | 100.00 | -48.40 | | CR | No | Yes |
| T3 | Nude | 72.13 | -59.97 | | PR | No | |
| T4 | Nude | 67.19 | -67.19 | | PR | Yes | No |
| T5 | Nude | 70.83 | -70.82 | | PR | Yes | Yes |
| T6 | Nude | 55.04 | 2.63 | | PR | No | |
| T7 | Nude | 68.50 | -61.79 | | PR | No | |
| T8 | Nude | 0.00 | 54.36 | 2.47 | PD2 | No | |
| T9 | Nude | 69.12 | -69.12 | | PR | Yes | No |
| T10 | Nude | 20.95 | -0.77 | | SD | No* | |
| T11 | Nude | 29.77 | -3.26 | | SD | No | |
| T12 | Nude | 62.30 | -62.30 | | PR | Yes | No |
| T13 | Nude | 61.63 | -61.63 | | PR | Yes | No |
| T14 | Nude | 44.47 | -44.47 | | SD | Yes | No |
| T15 | Nude | 63.83 | -63.83 | | PR | Yes | No |
| T16 | Nude | 63.14 | -63.14 | | PR | Yes | No |
| T17 | Nude | 100.00 | -100.00 | | CR | No | Yes |
| Cis 1 | Nude | 39.84 | 31.65 | 3.46 | PD2 | No | |
| Cis 2 | Nude | 0.00 | 211.32 | 2.61 | PD2 | No | |
| Cis 3 | Nude | 23.07 | 276.37 | 2.38 | PD2 | No | |
| Cis 4 | Nude | 20.68 | 217.08 | 2.61 | PD2 | No | |
| Cis 5 | Nude | 23.58 | 11.21 | | SD | No | |
| Cis 6 | Nude | 20.16 | -6.17 | | SD | No | |
| Cis 7 | Nude | 26.51 | 209.30 | 2.61 | PD2 | No | |
| Cis 8 | Nude | 27.34 | 48.96 | 3.31 | PD2 | No | |
| Cis 9 | Nude | 0.00 | 269.08 | 2.38 | PD2 | No | |
| Cis 10 | Nude | 0.00 | 317.82 | 1.54 | PD2 | No | |

*Euthanized early due to weight loss. Included in the PC group due to collection time point.

**Table S1. Individual PDX responses to COJEC or Cisplatin treatment.** Detailed data of the individual response to treatment following the parameters of The pediatric preclinical testing program *(50)*. Max regression (%) represents the maximum tumor volume reduction during treatment time. The end of treatment was day 40 or the day of surgery. T – COJEC-treated mice, HD – high dose, Cis – Cisplatin treated mice.
